# Human Galectin-9 Potently Enhances SARS-CoV-2 Replication and Inflammation in Airway Epithelial Cells

**DOI:** 10.1101/2022.03.18.484956

**Authors:** Li Du, Mohamed S. Bouzidi, Akshay Gala, Fred Deiter, Jean-Noël Billaud, Stephen T. Yeung, Prerna Dabral, Jing Jin, Graham Simmons, Zain Dossani, Toshiro Niki, Lishomwa C. Ndhlovu, John R. Greenland, Satish K. Pillai

**Affiliations:** Vitalant Research Institute, San Francisco, CA, USA; Department of Laboratory Medicine, University of California, San Francisco, CA, USA; Department of Medicine, University of California, San Francisco, CA, USA; Veterans Affairs Health Care System, San Francisco, CA, USA; QIAGEN Digital Insights, Redwood City, CA, USA; Division of Infectious Diseases, Department of Medicine, Weill Cornell Medicine, New York, NY, USA; Kagawa University, Kagawa, Japan

## Abstract

The severe acute respiratory syndrome coronavirus 2 (SARS-CoV-2) pandemic has caused a global economic and health crisis. Recently, plasma levels of galectin-9 (Gal-9), a β-galactoside-binding lectin involved in immune regulation and viral immunopathogenesis, were reported to be elevated in the setting of severe COVID-19 disease. However, the impact of Gal-9 on SARS-CoV-2 infection and immunopathology remained to be elucidated. Here, we demonstrate that Gal-9 treatment potently enhances SARS-CoV-2 replication in human airway epithelial cells (AECs), including primary AECs in air-liquid interface (ALI) culture. Gal-9-glycan interactions promote SARS-CoV-2 attachment and entry into AECs in an ACE2-dependent manner, enhancing the binding affinity of the viral spike protein to ACE2. Transcriptomic analysis revealed that Gal-9 and SARS-CoV-2 infection synergistically induce the expression of key pro-inflammatory programs in AECs including the IL-6, IL-8, IL-17, EIF2, and TNFα signaling pathways. Our findings suggest that manipulation of Gal-9 should be explored as a therapeutic strategy for SARS-CoV-2 infection.

**Importance:** COVID-19 continues to have a major global health and economic impact. Identifying host molecular determinants that modulate SARS-CoV-2 infectivity and pathology is a key step in discovering novel therapeutic approaches for COVID-19. Several recent studies have revealed that plasma concentrations of the human β-galactoside-binding protein galectin-9 (Gal-9) are highly elevated in COVID-19 patients. In this study, we investigated the impact of Gal-9 on SARS-CoV-2 pathogenesis *ex vivo* in airway epithelial cells (AECs), the critical initial targets of SARS-CoV-2 infection. Our findings reveal that Gal-9 potently enhances SARS-CoV-2 replication in AECs, interacting with glycans to enhance the binding between viral particles and entry receptors on the target cell surface. Moreover, we determined that Gal-9 accelerates and exacerbates several virus-induced pro-inflammatory programs in AECs that are established signature characteristics of COVID-19 disease and SARS-CoV-2-induced acute respiratory distress syndrome (ARDS). Our findings suggest that Gal-9 is a promising pharmacological target for COVID-19 therapies.

## Introduction

In December 2019, the first known case of coronavirus disease 2019 (COVID-19), caused by SARS-CoV-2 infection, was reported in Wuhan, China. Rapidly, COVID-19 cases were reported worldwide. To date, SARS-CoV-2 has accounted for more than 360 million infections and more than 5.6 million deaths worldwide (1). The rapid spread of SARS-CoV-2 continues to have a major impact on global health and the economy. COVID-19 is mainly characterized by pneumonia, including fever, cough, and chest discomfort, and in severe cases dyspnea and lung infiltration (2).

The major cause of death in COVID-19 cases is acute respiratory distress syndrome (ARDS) accompanied by a cytokine storm (3). Several reports have identified specific circulating proteins and cytokines in blood plasma that are elevated in the setting of COVID-19 and may constitute clinically useful disease biomarkers (4, 5). One such specific protein is Gal-9. Studies have recently revealed that plasma Gal-9 levels are elevated in COVID-19 patients and are positively correlated with COVID-19 severity (6–9). Furthermore, plasma levels of Gal-9 during COVID-19 were positively correlated with key pro-inflammatory cytokines, including interleukin 6 (IL-6), interferon gamma-induced protein 10 (IP-10), and tumor necrosis factor alpha (TNFα) (6). However, the mechanism linking Gal-9 to severe COVID-19 disease remains to be elucidated.

Gal-9 belongs to the galectin family, which includes 15 carbohydrate binding proteins sharing a common carbohydrate-recognition domain (CRD) (10). Conserved CRDs of galectins can homodimerize and have strong binding affinity for poly-N-acetyllactosamine (poly-LacNAc), which is present on the branches of *N*- and *O*-linked glycans (11). Gal-9 is known for its regulation of immune responses and viral pathogenesis through glycan-mediated recognition. It is ubiquitously expressed in different tissues and cells in humans (e.g. endothelial cells, T lymphocytes, dendritic cells (DCs), macrophages, intestinal epithelial cells) and is localized in the extracellular matrix, surface, cytoplasm and nucleus of cells (12). Circulating levels of Gal-9 serve as sensitive and non-invasive biomarkers in a broad range of conditions including cancer, autoimmunity, and infectious diseases, and the roles of Gal-9 vary with respect to cell type and disease state (13). For example, Gal-9 suppresses antigen-specific CD8+ T cell effector functions via interaction with its receptor, TIM-3 (14), and the Gal-9/TIM-3 axis promotes tumor survival through cross-talk with the PD-1 immune checkpoint (15). The functions of Gal-9 in the inflammatory response have been studied extensively. Gal-9 enhances cytokine secretion in the human mast cell line (16), and potentiates secretion of pro-inflammatory cytokines in inflammatory models of arthritis via TIM-3 interaction (17). In specific relevance to viral pathogenesis, Gal-9 can bind to glycan structures expressed on the surface of both host cells and microorganisms to modulate antiviral immunity, and to promote or inhibit viral infection and replication (18). Gal-9 was demonstrated to inhibit human cytomegalovirus (HCMV) infection by its carbohydrate recognition domains (18). In hepatitis C virus (HCV)-infected individuals, virus infection induces Gal-9 secretion, which in turn induces pro-inflammatory cytokines leading to depletion of CD4+ T cells, apoptosis of HCV-specific cytotoxic T cells (CTLs), and expansion of regulatory T cells (Tregs) (19). Gal-9 is elevated in human immunodeficiency virus-1 (HIV-1)-infected individuals (20, 21), mediates HIV-1 transcription and reactivation (20, 22), and potentiates HIV-1 infection by regulating the T cell surface redox environment (23). These previous reports provide compelling evidence of the diverse roles of Gal-9 in viral infection and virus-associated immunopathology.

To date, a causal role of Gal-9 in SARS-CoV-2 pathology has not been demonstrated. To address this gap, we investigated the effects of Gal-9 treatment on SARS-CoV-2 replication and pro-inflammatory signaling in immortalized and primary human airway epithelial cells (AECs). Our data show that Gal-9 facilitates SARS-CoV-2 replication and promotes virus-associated immunopathology in the human airway, motivating exploration into Gal-9 manipulation as a therapeutic strategy for COVID-19 disease.

## Results

### Gal-9 potently enhances SARS-CoV-2 replication in Calu-3 cells

To investigate the impact of Gal-9 on SARS-CoV-2 infection, we first determined the cytotoxicity of recombinant stable-form Gal-9, to optimize dosing in the immortalized Calu-3 AEC line. The 50% cytotoxic concentration (CC_50_) value detected by MTT assay was 597 nM for Gal-9 in Calu-3 cells (Fig. 1A). Next, we evaluated the effects of Gal-9 on SARS-CoV-2 replication at 50 nM, 100 nM, and 250 nM concentrations. Calu-3 cells were pre-treated with Gal-9 for six hours before viral infection (MOI=0.01), and Gal-9 was maintained in the media until 24 hours following infection. SARS-CoV-2 infection, as measured by quantitation of viral nucleocapsid (*N*) gene expression, was increased significantly by treatment with Gal-9 in a dose dependent manner (*p<0.0001*) (Fig. 1B), with up to 27-fold induction at the highest concentration of Gal-9. Similarly, release of infectious virus in the supernatant was enhanced significantly by Gal-9 in a dose dependent manner, as measured by median tissue culture infectious dose (TCID_50_) (*p<0.05*) (Fig. 1C). The enhancement of virus production by Gal-9 was confirmed by specific staining of viral nucleocapsid protein using an immunofluorescence assay (IFA) (Fig. 1D,E). Taken together, these data demonstrate that Gal-9 increases SARS-CoV-2 viral production in susceptible Calu-3 cells.

**Fig. 1:**
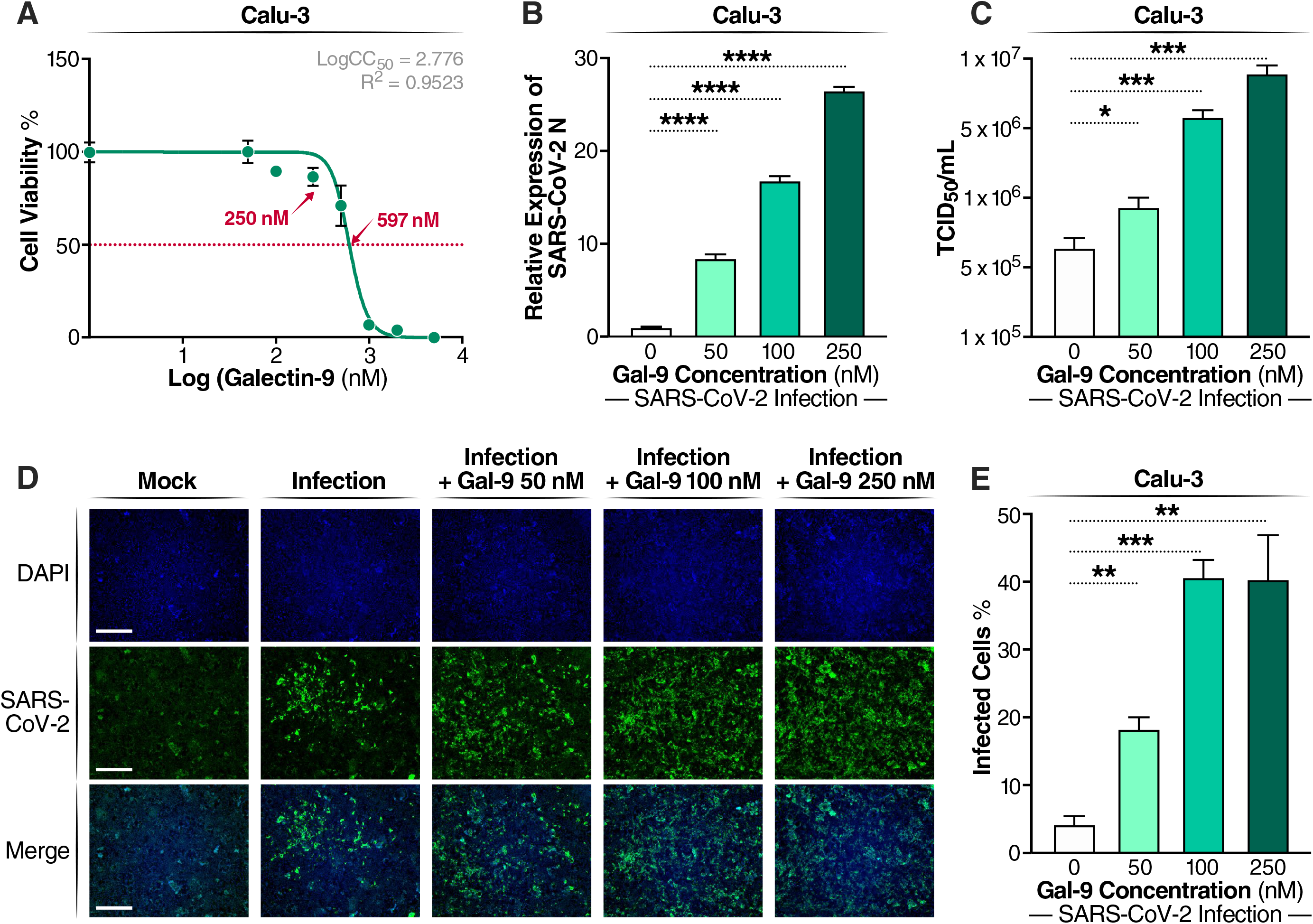
Gal-9 increases virus production in SARS-CoV-2-infected Calu-3 cells. (A) Cellular toxicity was examined in Calu-3 cells using an MTT assay and was expressed as relative cell viability as compared to Gal-9-untreated control (set at 100%). The LogCC_50_ value for Gal-9 is displayed. The red arrows represent 250 nM and the CC_50_ value (597 nM) of Gal-9, respectively. (B) The effect of Gal-9 on viral *N* gene expression in Calu-3 cells was measured by RT-qPCR. Cells were pretreated with Gal-9 at the indicated concentrations for six hours, followed by infection with SARS-CoV-2 (MOI=0.01) for 24 h in the presence of Gal-9. 24 hpi, cells were collected for RNA isolation and RT-qPCR targeting the *N* gene. (C) Infectious virus release in the supernatant of SARS-CoV-2-infected Calu-3 cells treated with varying doses of Gal-9 as described in (B) were measured using TCID_50_. (D) Immunofluorescence staining of Calu-3 cells with DAPI (blue) or anti-N Ab (green) pretreated with Gal-9 at the indicated concentrations, followed by infection with SARS-CoV-2 as described in (B). Scale bar, 500 μM. (E) Quantification of SARS-CoV-2 infected cells in Calu-3 cells (shown in panel D). Data are representative of the results of three independent experiments (mean ± SEM). Statistical significance was analyzed by *t* test. *p*_≤_*0.05* [*], *p*_≤_ *0.01* [**], *p*_≤_*0.001* [***], *p*_≤_*0.0001* [****].

### Gal-9 enhances SARS-CoV-2 entry in an ACE2-dependent manner

To investigate the stage of the virus replication cycle impacted by Gal-9, we treated Calu-3 cells with Gal-9 at the concentration of 250 nM, chosen based on our analyses of toxicity and dose response. Calu-3 cells were treated with Gal-9 before and after virus infection. Protocols are illustrated in Extended Data Fig. 1A. Gal-9-mediated enhancement of virus production was significantly higher in pre-treated cells (*p<0.05*), as compared to cells treated with Gal-9 following viral exposure (Extended Data Fig. 1B). These results suggested that Gal-9 likely impacts the early stage of the SARS-CoV-2 viral life cycle.

Based on the results of our time-course experiments, we sought to determine whether Gal-9 affects SARS-CoV-2 viral entry. Firstly, we examined the role of Gal-9 in SARS-CoV-2 cell-surface attachment. Cells were incubated with SARS-CoV-2 at 4°C for 2 hours and attached SARS-CoV-2 viral particles were detected after washing the cells three times. Gal-9 induced a substantial, highly significant (*p<0.0001*), and dose-dependent increase in cell-surface SARS-CoV-2 attachment (Fig. 2A). We next determined the capacity of Gal-9 to affect the entry of VSV-SARS-CoV-2 spike-ΔG-luciferase reporter pseudovirus (SARS-2-S) into Calu-3 cells. Positive serum (P Serum), which was predetermined to possess SARS-CoV-2 neutralizing activity, potently reduced SARS-2-S infection (*p<0.01*) but did not suppress VSV-G infection (Fig. 2B). Gal-9 markedly enhanced SARS-2-S infection in a dose-dependent manner in Calu-3 cells which endogenously express ACE2 and TMPRSS2 (*p<0.01*) (Fig. 2B), indicating that Gal-9 can potentiate SARS-CoV-2 attachment and entry. Unexpectedly, entry of VSV-G was also significantly enhanced by Gal-9 treatment (*p<0.0001*), suggesting that this pro-viral activity may be generalizable to other viral taxa.

**Fig. 2:**
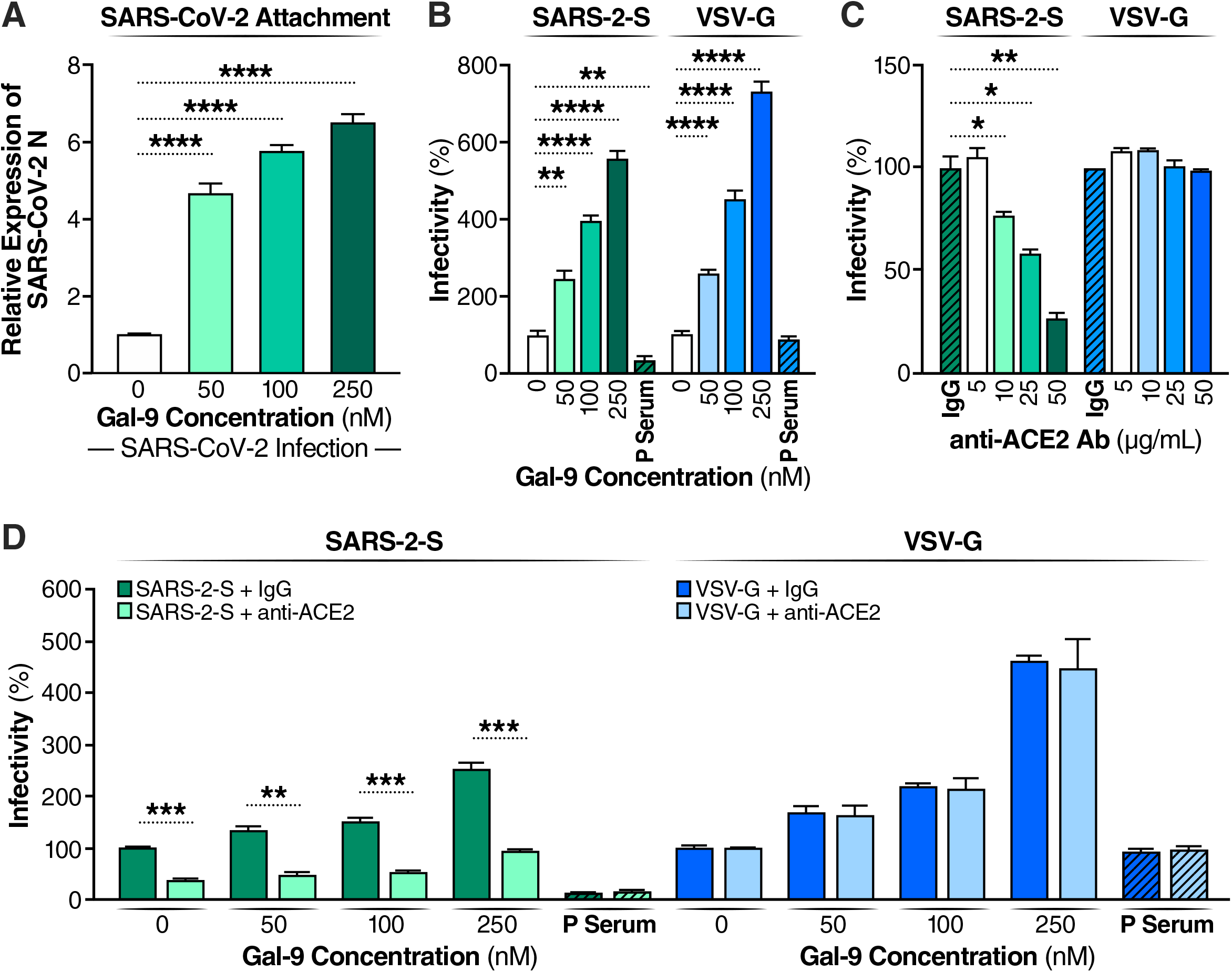
Gal-9 facilitates the cellular attachment and entry of SARS-CoV-2. (A) Attached SARS-CoV-2 virions on the cell surface were detected by RT-qPCR. Calu-3 cells were pretreated with Gal-9 for one hour, then cells were incubated with SARS-CoV-2 (MOI=0.01) in solutions with or without Gal-9 at the indicated concentrations at 4°C for two hours. Cells were washed three times with PBS and harvested for RNA isolation and RT-qPCR measurement of SARS-CoV-2 *N* gene expression. (B) Relative infectivity of SARS-2-S pseudotyped virus and VSV-G pseudotyped virus in Calu-3 treated with Gal-9 at the indicated concentrations. Calu-3 cells were exposed to Gal-9 for six hours and then infected with SARS-2-S pseudotyped virus or VSV-G pseudotyped virus in solutions containing Gal-9 at the indicated concentrations. Pseudotyped viral entry was analyzed by luciferase activity 24 hpi. Positive serum predetermined to possess anti-SARS-CoV-2 neutralizing activity was used as a negative control. Luciferase signals obtained in the absence of Gal-9 were used for normalization. (C) Relative infectivity of SARS-2-S pseudotyped virus and VSV-G pseudotyped virus in Calu-3 cells treated with anti-ACE2 Ab at the indicated concentrations. Calu-3 cells pre-incubated with anti-ACE2 Ab (R&D Systems, AF933) at the indicated concentrations, or control antibody (anti-goat IgG (R&D Systems, AB-108-C), 50 μg/ml) were co-administered with SARS-2-S pseudotyped and VSV-G pseudotyped virus. At 24 hpi, pseudotyped viral entry was analyzed by luciferase activity. Luciferase signals obtained using the control antibody (anti-goat IgG, 50 μg/ml) were used for normalization. (D) The effect of anti-ACE2 Ab on Gal-9-enhanced cell entry of SARS-2-S was evaluated by measuring luciferase activity. Calu-3 cells were pretreated with Gal-9 for six hours and then pre-incubated with anti-ACE2 Ab (25 μg/ml) or control antibody for one hour, and cells were inoculated with SARS-2-S pseudotyped or VSV-G pseudotyped virus in a solution containing Gal-9 at the indicated concentrations. At 24 hpi, pseudotyped viral entry was analyzed by luciferase activity. Luciferase signals obtained in the absence of both Gal-9 and anti-ACE2 Ab were used for normalization. Data are representative of the results of three independent experiments (mean ± SEM). Statistical significance was analyzed by *t* test. *p*_≤_*0.05* [*], *p*_≤_ *0.01* [**], *p*_≤_*0.001* [***], *p*_≤_*0.0001* [****].

ACE2 has been identified as the critical receptor for SARS-CoV-2 binding and entry (24). To explore whether Gal-9-promoted virus entry depends on ACE2 binding, we treated cells with an anti-ACE2 antibody that competitively binds to the receptor. As expected, anti-ACE2 antibody blocked SARS-2-S but not VSV-G entry in a dose dependent manner (*p<0.05*) (Fig. 2C). Anti-ACE2 antibody also blocked Gal-9-enhanced virus infection (*p<0.01*) (Fig. 2D), indicating that Gal-9 facilitates SARS-CoV-2 entry in an ACE2-dependent manner.

We next investigated the potential mechanisms underlying the Gal-9-mediated enhancement of SARS-CoV-2 viral entry. SARS-CoV-2 can use the endosomal cysteine proteases cathepsin B and L (CatB/L) and the serine protease TMPRSS2 to prime entry (25). Only TMPRSS2 activity is essential for viral spread and pathogenesis in the infected host, whereas CatB/L activity is dispensable. In Calu-3 cells, which express ACE2 and TMPRSS2 (Fig. 3A), SARS-CoV-2 entry was demonstrated to be primed by TMPRSS2 (26). To determine if Gal-9 modulates ACE2 and TMPRSS2 surface expression to promote virus entry, we measured Calu-3 cell surface ACE2 and TMPRSS2 expression by flow cytometry. Gal-9 exhibited no effects on ACE2 and TMPRSS2 expression (Fig. 3B), suggesting that Gal-9 facilitates virus entry independently of ACE2 and TMPRSS2 induction. We next investigated the direct impact of Gal-9 on the interaction between the SARS-CoV-2 spike protein and the ACE2 entry receptor, using purified spike and ACE2 proteins in an established sandwich ELISA protocol (27). Gal-9 significantly enhanced binding between ACE2 and spike (*p<0.05*) (Fig. 3C), indicating Gal-9 facilitates virus entry, and viral replication at large, by strengthening ACE2 and spike interaction.

**Fig. 3:**
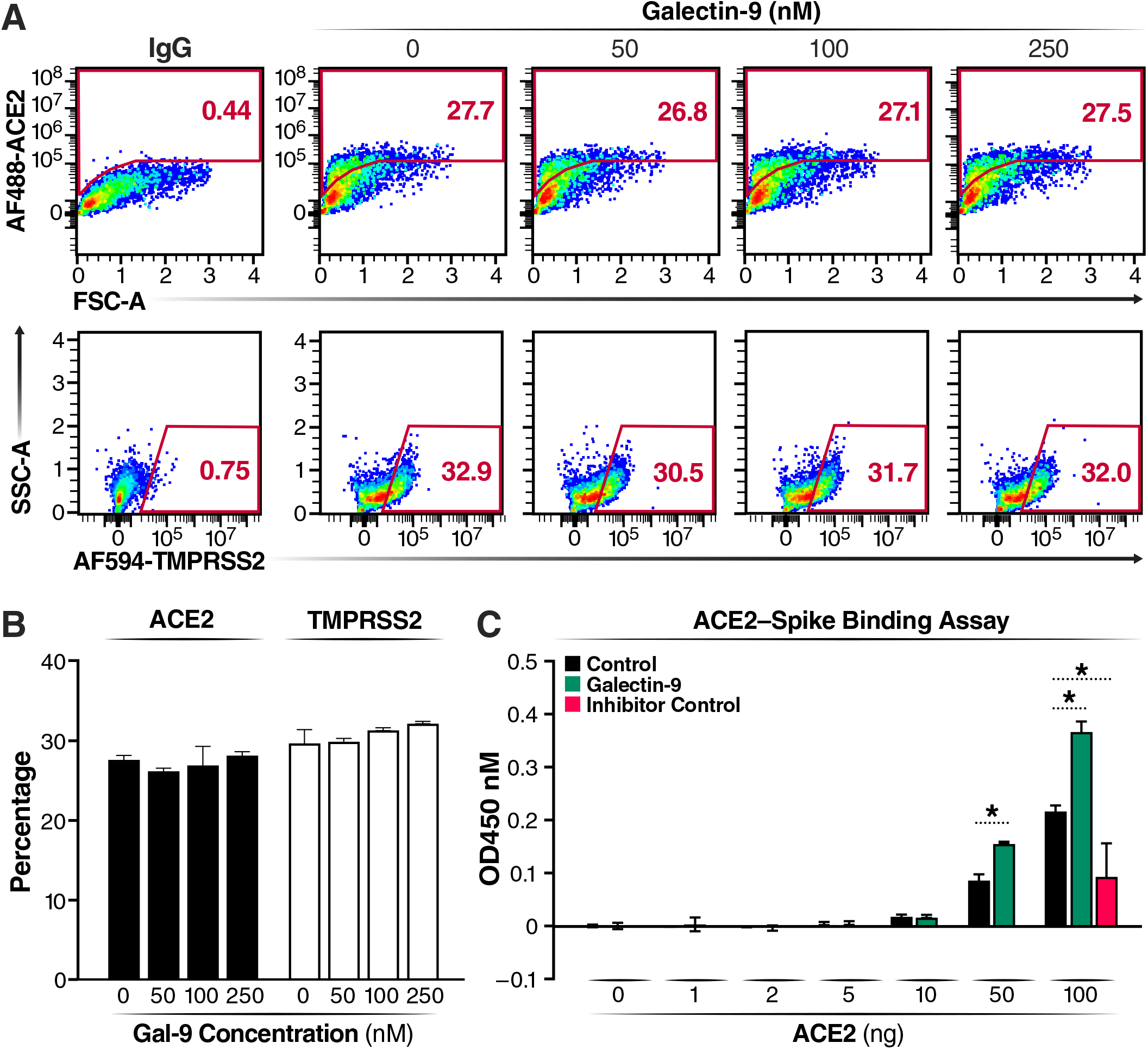
Gal-9 enhances the binding of SARS-CoV-2 spike to ACE2 without affecting ACE2 and TMPRSS2 cell surface expression. (A) Representative flow cytometry plot describing the protein levels of ACE2 and TMPRSS2 on the surface of Calu-3 cells treated with Gal-9. Calu-3 cells were treated with Gal-9 at the indicated concentrations for 24 h. Cells were then washed and detached before antibody staining for flow cytometry. (B) Percentages of cells expressing ACE2 or TMPRSS2 at the cell surface, measured using flow cytometry. Data represent the results of three independent experiments (mean ± SEM). (C) The dose-response of SARS-CoV-2 spike-ACE2 binding activity measured by reading the absorbance at the wavelength of 450 nm. Data are representative of the results of three independent experiments (mean ± SEM). Statistical significance was analyzed by *t* test. *p*_≤_*0.05* [*], *p*_≤_ *0.01* [**], *p*_≤_*0.001* [***], *p*_≤_*0.0001* [****].

### Gal-9-mediated enhancement of SARS-CoV-2 entry is glycan-dependent

To explore the impact of Gal-9-glycan interactions on SARS-CoV-2 infection, we first determined if Gal-9-enhanced SARS-CoV-2 entry was dependent on CRD activity, relying on lactose, a competitive inhibitor of galectin carbohydrate-binding activity (28). Our data demonstrated that lactose treatment significantly abrogated Gal-9-mediated SARS-CoV-2 entry in a dose-dependent manner (*p<0.001*) (Fig. 4A). We then deglycosylated Calu-3 target cells using two complementary approaches, kifunensine treatment and PNGase F exposure (29). Kifunensine inhibits mannosidase I enzymatic activity within the cell, preventing hybrid and complex N-linked glycosylation of synthesized proteins, and PNGase F chemically removes N glycans from the cell surface. Loss of host complex N-glycans achieved using both of these strategies led to significant inhibition of Gal-9-enhanced SARS-CoV-2 entry (*p<0.01*) (Fig. 4B,C). In the absence of Gal-9 treatment, PNGase F administration inhibited SARS-CoV-2 entry (*p<0.05*) (Fig. 4C). Collectively, these data demonstrate that Gal-9 promotes SARS-CoV-2 attachment and entry into host cells in a glycan-dependent manner.

**Fig. 4:**
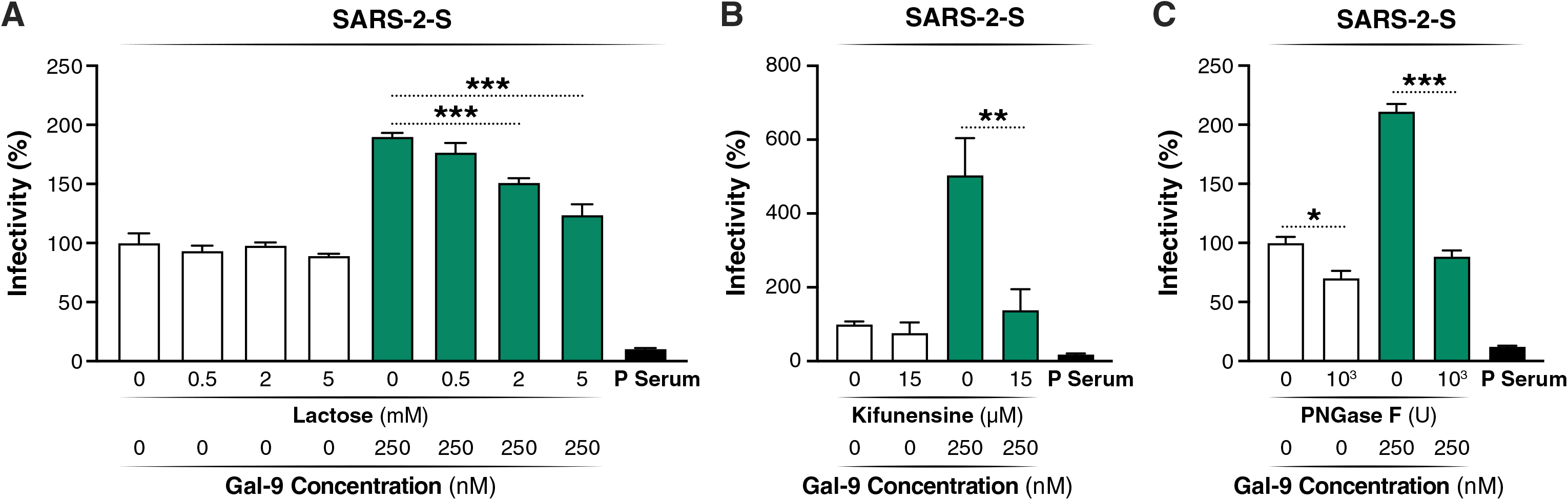
Gal-9 mediated enhancement of SARS-CoV-2 entry is glycan dependent. (A) The effect of lactose on Gal-9-enhanced SARS-CoV-2 infection was evaluated by measuring luciferase activity. Calu-3 cells were pre-treated with Gal-9 and lactose at the indicated concentrations for six hours. Cells were then inoculated with SARS-2-S pseudotyped virus, maintaining Gal-9 and lactose at indicated concentrations. Pseudotyped viral entry was analyzed by luciferase activity at 24 hpi. Luciferase signals obtained in the absence of Gal-9 and lactose were used for normalization. (B) The effect of kifunensine on Gal-9-enhanced SARS-CoV-2 entry was detected by measuring luciferase activity. Calu-3 cells were pre-incubated with kifunensine for 24 hours. Cells were then treated with Gal-9 and kifunensine for six hours. Cells were then inoculated with SARS-2-S pseudotyped virus in a solution containing Gal-9 and kifunensine at the indicated concentrations. Pseudotyped viral entry was analyzed by luciferase activity at 24 hpi. Positive serum (P Serum) predetermined to possess anti-SARS-CoV-2 neutralizing activity was used as a negative control. Luciferase signals obtained in the absence of Gal-9 and kifunensine were used for normalization. (C) The effect of PNGase F on Gal-9-enhanced SARS-CoV-2 entry was detected by measuring luciferase activity. Calu-3 cells were pre-incubated with PNGase F for four hours. Cells were then treated with Gal-9 for six hours, and were then inoculated with SARS-2-S pseudotyped virus in a solution containing Gal-9. Pseudotyped viral entry was analyzed by luciferase activity at 24 hpi. Luciferase signals obtained in the absence of Gal-9 and PNGase F were used for normalization. Data are representative of the results of three independent experiments (mean ± SEM). Statistical significance was analyzed by *t* test. *p*_≤_*0.05* [*], *p*_≤_ *0.01* [**], *p*_≤_*0.001* [***], *p*_≤_*0.0001* [****].

### Gal-9 promotes SARS-CoV-2-associated expression of pro-inflammatory cytokines

We next evaluated the temporal characteristics of Gal-9-mediated enhancement of SARS-CoV-2 replication. Our growth curves are compatible with the concept that Gal-9 facilitates SARS-CoV-2 entry, as the expression of the viral *N* gene was significantly increased within one to three hours post-infection (hpi) in Calu-3 cells (*p<0.0001*), as compared to untreated controls (Fig. 5A). Maximum viral yields were detected at 36 hpi with Gal-9 treatment, and 48 hpi without Gal-9 treatment. In accordance with the viral growth kinetic data, microscopy images showed that virus-mediated cytopathic effects (CPE) were much more pronounced in infected cultures treated with Gal-9 at both 36 and 72 hpi, as compared to infected, Gal-9 untreated cultures (Fig. 5B). The enhancement of virus production by Gal-9 over time was also evaluated by TCID_50_ in the supernatant at various time points (Fig. 5C). Enhanced release of infectious virus in the presence of Gal-9 was observed as early as nine hpi (*p<0.05*), again reflecting effects of Gal-9 on early stages of the viral life cycle. Collectively, these results reinforce the concept that Gal-9 promotes SARS-CoV-2 viral production through enhancement of viral entry.

**Fig. 5:**
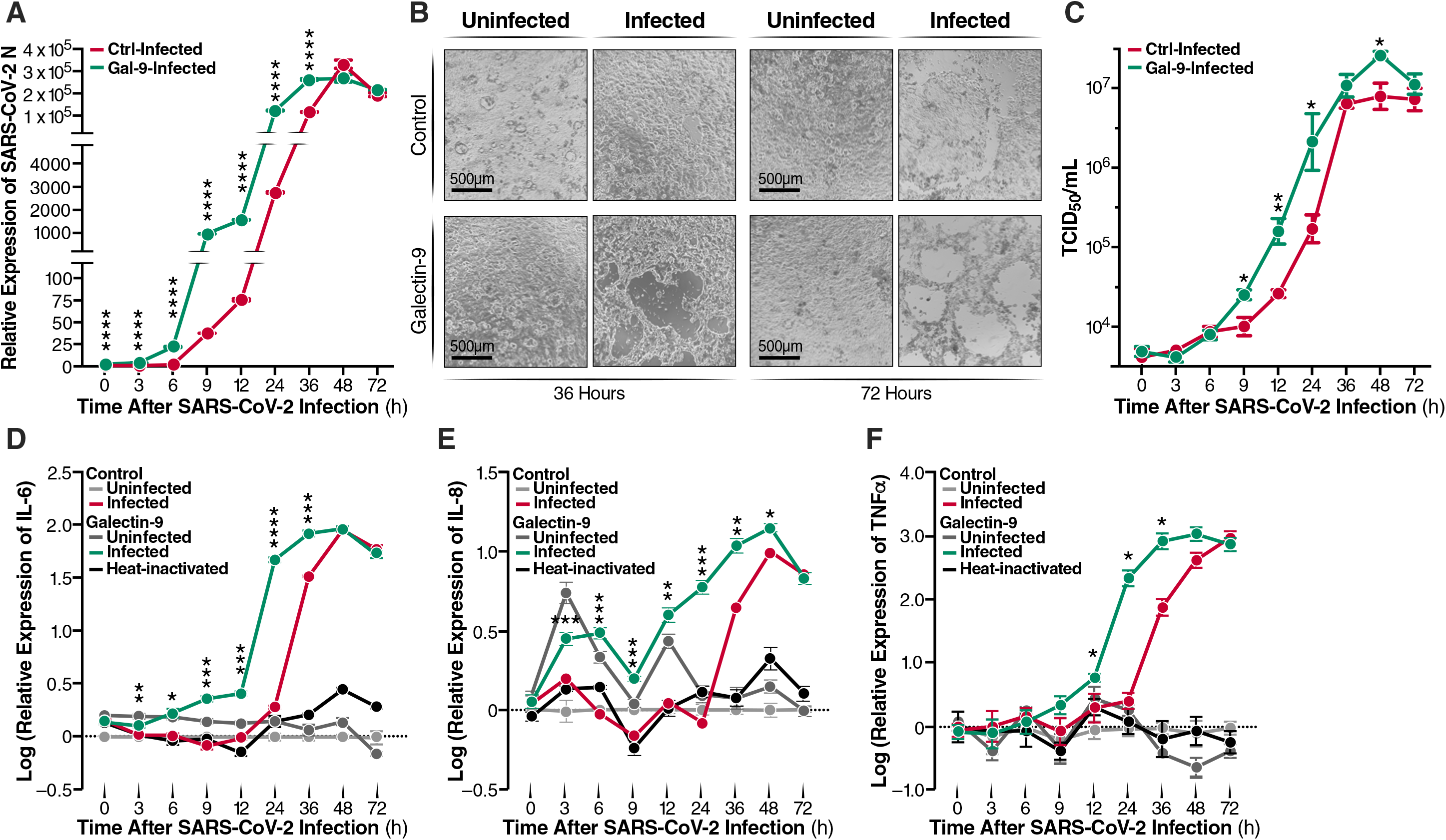
Gal-9 accelerates and increases SARS-CoV-2-mediated inflammatory response. (A) SARS-CoV-2 growth curves in Calu-3 cells pre-treated with or without Gal-9 (250 nM) for six hours and then infected with SARS-CoV-2 (MOI=0.01) in the presence or absence of Gal-9. Cells were collected at the indicated times for RNA isolation and RT-qPCR targeting *N* gene. (B) Calu-3 cells from panel A were observed for development of CPE by bright-field microscopy at 36 hpi and 72 hpi. Scale bar, 500 μM. (C) Viral titer (TCID_50_/ml) of supernatant collected at the indicated time points after SARS-CoV-2 infection of Gal-9 treated or untreated Calu-3 cells as described in (A). (D-F) mRNA levels of pro-inflammatory cytokines *IL-6* (D), *IL-8* (E), and *TNF*_α_ (F) were detected by RT-qPCR at the indicated time points. Heat-inactivated = infected with heat-inactivated SARS-CoV-2. Data are representative of the results of three independent experiments (mean ± SEM). Statistical significance was analyzed by *t* test. *p*_≤_*0.05* [*], *p*_≤_ *0.01* [**], *p*_≤_*0.001* [***], *p*_≤_*0.0001* [****].

The host inflammatory response drives much of the pathology, morbidity, and mortality associated with SARS-CoV-2 infection (30), and Gal-9 was positively correlated with pro-inflammatory mediators and disease severity in COVID-19 patients. Therefore, we sought to determine if Gal-9 promotes pro-inflammatory cytokine expression, either directly, or indirectly via enhancement of viral replication. Uninfected, untreated cells were also characterized as a negative control and reference. In the absence of SARS-CoV-2 infection, Gal-9 treatment induced *IL-6*, *IL-8*, and *TNF*_α_ (all *p<0.05*) gene expression in Calu-3 cells, albeit at modest levels (Fig. 5D-5F). SARS-CoV-2 infection potently induced *IL-6*, *IL-8*, and *TNF*_α_ (all *p<0.05*), starting at 24, 36, and 24 hpi, respectively (Fig. 5D-5F). In the presence of Gal-9, SARS-CoV-2 infection significantly induced *IL-6*, *IL-8*, and *TNF*_α_ (all *p<0.05*), starting at 9, 12, and 9 hpi, respectively (Fig. 5D-5F). Gal-9 treatment of infected cultures potentiated *IL-6* (*p<0.001*), *IL-8* (*p<0.05*), and *TNF*_α_ (*p<0.05*) expression as compared to SARS-CoV-2 infection alone (Fig. 5D-5F). Moreover, Gal-9 treatment of infected cultures significantly increased *IL-6*, *IL-8*, and *TNF*_α_ (all *p<0.05*) expression as compared to Gal-9 treatment in the absence of SARS-CoV-2 infection. These observations were confirmed, in part, at the protein level by applying a bead-based immunoassay to culture supernatants. Protein measurements confirmed enhanced IL-6 (*p<0.05*) and IL-8 secretion (*p<0.01*) by Gal-9 in the presence of virus (Extended Data Fig. 2A,B). TNFα and IL-17A secretion were additionally induced by Gal-9 in infected cells, although not quite achieving statistical significance (Extended Data Fig. 2C,D). Considering that the effects of Gal-9 on cytokine expression were relatively negligible in the absence of infection, our data indicate that Gal-9 and SARS-CoV-2 synergistically promote expression of pro-inflammatory cytokines in AECs. A formal quantitative analysis of synergy indeed revealed a synergistic impact of combined Gal-9 exposure and SARS-CoV-2 infection on *IL-6* and *TNF*_α_ expression in AECs (Extended Data Fig. 3).

### RNA-seq analysis reveals synergistic effects of Gal-9 and SARS-CoV-2 on the host transcriptome

To understand the transcriptional impact of Gal-9-mediated enhancement of SARS-CoV-2 infection and immunopathology, we performed RNA-seq analysis on Calu-3 cells infected for 24 hours with SARS-CoV-2 in the presence or absence of 250 nM Gal-9. Uninfected, untreated cells were also characterized as a negative control and reference for the other three experimental conditions. Differentially-expressed gene (DEG) analysis was performed, using a false discovery rate (FDR) cutoff of 0.05. Only one protein coding gene (RNU12) was significantly modulated by SARS-CoV-2 infection alone (FDR=1.89E-07) (Fig. 6A). 87 genes were significantly modulated by Gal-9 treatment alone (Fig. 6B), including 30 down-regulated and 57 up-regulated genes. Cells infected in the presence of Gal-9 exhibited a dramatic impact on the host transcriptome, with 1094 DEGs identified (Fig. 6C), including 323 down-regulated and 771 up-regulated genes. Using Ingenuity Pathway Analysis (IPA), we analyzed the enriched canonical pathways that overlap with the DEGs (Fig. 6D). Key pro-inflammatory programs including IL-17, EIF2, IL-6, IL-8, and JAK/STAT signaling pathways were activated in infected cells in the presence of Gal-9. A detailed interactome depicting Gal-9 and SARS-CoV-2 effects on IL-17 and IL-6 signaling pathway members are shown in Extended Data Fig. 4 and Extended Data Fig. 5.

**Fig. 6:**
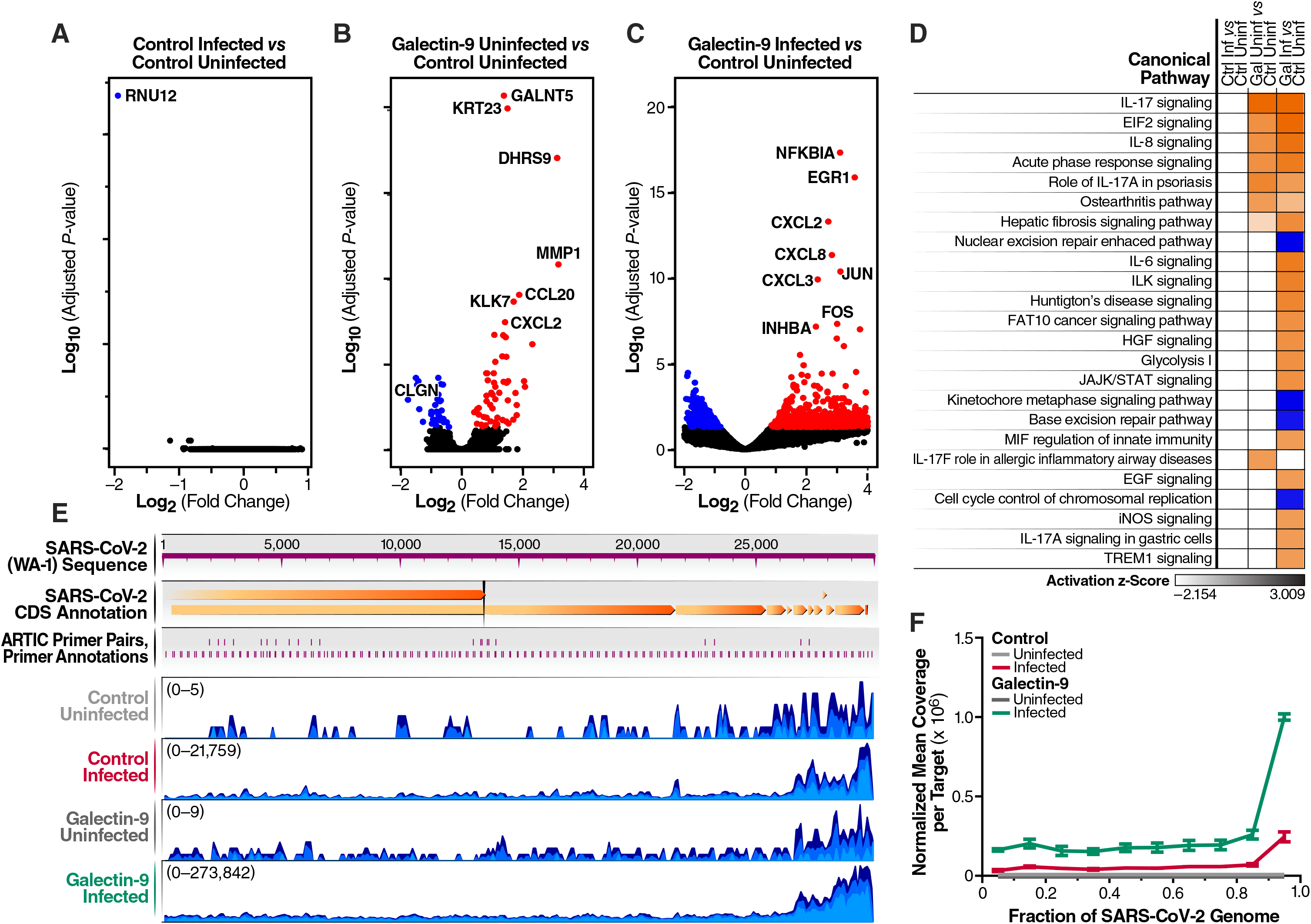
Impact of Gal-9 treatment on the transcriptome of SARS-CoV-2-infected Calu-3 cells. (A-C) Volcano plots showing the proportion of differentially-expressed genes (DEGs) in the setting of SARS-CoV-2 infection (Control Infected vs Control Uninfected) (A), Gal-9 treatment (Galectin-9 Uninfected vs Control Uninfected) (B), and SARS-CoV-2 infection in the presence of Gal-9 (Galectin-9 Infected vs Control Uninfected) (C). DEGs (FDR<0.05) with log2(fold change) > 0 are indicated in red. DEGs (FDR<0.05) with log2(fold change) < 0 are indicated in blue. Non-significant DEGs are indicated in black. Ctrl = Control. Gal = Galectin-9. Inf = Infected with SARS-CoV-2. Uninf = Uninfected with SARS-CoV-2. (D) Top enriched canonical pathways identified using IPA. The orange and blue-colored bars in the bar chart indicate predicted pathway activation or predicted inhibition, respectively, based on z-score. (E) Sample coverage tracks from the QIAGEN genome browser depicting SARS-CoV-2 assembly. Sample coverage tracks were obtained by mapping raw sequencing reads to the USA-WA1/2020 reference genome. Mapped read counts of Control Uninfected, Control Infected, Galectin-9 Uninfected, and Galectin-9 Infected are 0 to 5, 0 to 21759, 0 to 9, and 0 to 273842, respectively. (F) Normalized mean coverage per targeted region of the SARS-CoV-2 genome. The X axis represents the relative target region positions of the SARS-CoV-2 genome. Data were normalized to the total read depth for each sample.

DEGs were also categorized with respect to diseases and host functions. Diseases and functions are listed in Extended Data Fig. 6A and B. The results demonstrate that inflammatory response, infectious, inflammatory, and immunological disease pathways were activated in the setting of Gal-9 treatment. Interestingly, respiratory disease and RNA post-transcriptional modification were specifically activated in cells infected with SARS-CoV-2 in the presence of Gal-9.

We next leveraged the RNA-seq data to examine the impact of Gal-9 treatment on SARS-CoV-2 expression, achieved by aligning sequencing reads against the USA-WA1/2020 reference genome. The number of reads mapping to each region of the viral genome was calculated and interpreted to infer viral expression patterns (Fig. 6E). The transcription of SARS-CoV-2 exhibited an uneven pattern of expression along the genome, typically with a minimum depth in the first coding regions with ORFs 1a and 1b, and the maximum towards the 3’ end. This skewing likely reflects the relative abundance of these sequences due to the nested transcription of SARS-CoV-2 subgenomic RNAs; all viral transcript variants include the terminal 3’ genome segment (31). Importantly, in accordance with our quantitative PCR data, Gal-9 treatment increased the expression of SARS-CoV-2, resulting in a ∼10-fold increase in the number of sequencing reads mapping to the USA-WA1/2020 reference (Fig. 6E,F), maintaining and even amplifying the observed 3’ skewing of viral transcripts.

### Gal-9 enhances SARS-CoV-2 replication and increases TNF**α** expression in primary airway epithelial cells

Finally, we sought to investigate the effect of Gal-9 treatment on SARS-CoV-2 infection and inflammatory response in primary human airway epithelial cells cultured at the air/liquid interface (ALI). Recent studies have reported that human airway epithelial cells represent the primary gateway for SARS-CoV-2 infection upon colonization of a new host (32). Rapid viral replication in these cells lead to the release of pro-inflammatory cytokines, which causes airway damage and diminished patient survival. Thus, our ALI-cultured primary airway epithelial system is useful in modeling the *in vivo* effects of Gal-9 on SARS-CoV-2 infection *ex vivo*. We pretreated primary AECs from five healthy donors with Gal-9 and then infected them with the SARS-CoV-2 Gamma variant (Pango lineage designation P.1) for 36 hours. The Gamma variant was selected based on our observations demonstrating that the Gamma variant infects primary AECs more efficiently than the founder virus lineage (USA-WA1/2020) (data not shown). Productive infection was observed in all five donor cultures, based on expression of the SARS-CoV-2 *N* gene (Fig. 7A). Validating our observations in Calu-3 cells, Gal-9 treatment significantly increased SARS-CoV-2 replication (*p<0.05*), by up to 2.6-fold (Fig. 7A). We also assessed pro-inflammatory signatures by RT-PCR. Our results showed that SARS-CoV-2 significantly induced *TNF*_α_ as compared to mock control (*p<0.05*) (Fig. 7B). The mRNA levels of *IL-6* and *IL-8* were not affected by SARS-CoV-2 infection at 36 hpi. Gal-9 treatment significantly increased *IL-6* expression in the presence of virus (*p<0.05*), and induction of *TNF*_α_ was observed but failed to achieve statistical significance (*p=0.055*) (Fig. 7B). Taken together, our findings in primary AECs validate and extend our previous observations, confirming that Gal-9 promotes SARS-CoV-2 replication and associated pro-inflammatory signaling in the airway epithelium.

**Fig. 7:**
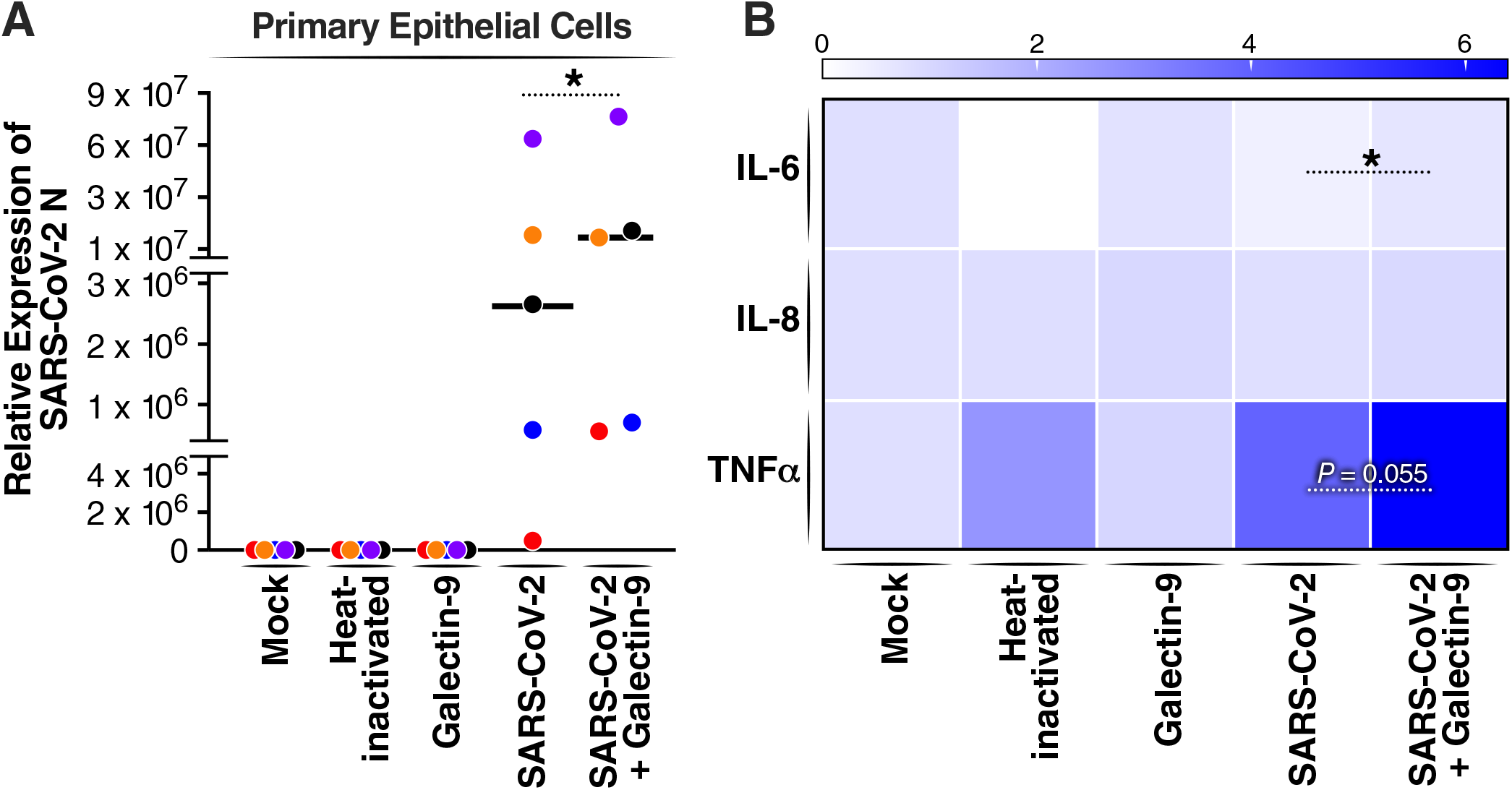
Gal-9 enhances SARS-CoV-2 replication and virus-induced inflammatory response in human primary airway epithelial cells. (A) RT-qPCR analysis of the SARS-CoV-2 *N* gene from ALI-cultured primary airway epithelial cells pre-treated with or without Gal-9 and infected with or without SARS-CoV-2 in five different donors. ALI-cultured primary epithelial cells were pre-incubated with or without Gal-9 (250 nM) for six hours and were then infected with SARS-CoV-2 (lineage P.1, MOI=0.1), heat-inactivated virus, or mock control, in a solution of Gal-9 (250 nM) at the apical chamber of inserts (250 μl) and the basal compartment (500 μl) for 2 h. Cells were washed and supplemented with fresh media alone or fresh media containing Gal-9 (250 nM). 36 hpi, cells were collected for RNA isolation and RT-qPCR. Each color represents data from one donor. (B) The mRNA level of pro-inflammatory cytokines *IL-6*, *IL-8*, and *TNF*_α_ were measured by RT-qPCR in ALI-cultured primary epithelial cells treated as described in (A) in five different donors. Data represent duplicate experiments for each donor (median). Statistical significance was analyzed by paired one-tailed Wilcoxon tests. *p*_≤_*0.05* [*], *p*_≤_ *0.01* [**], *p*_≤_*0.001* [***], *p*_≤_*0.0001* [****].

## Discussion

The elevation of plasma Gal-9 levels in COVID-19 cases and severe COVID-19 disease has been confirmed in multiple reports (6, 33). Here, we leveraged a recombinant stable-form of Gal-9 as a proxy for endogenously produced Gal-9, and investigated its impact on SARS-CoV-2 replication and host immune signaling. Our data reveal that Gal-9 enhances SARS-CoV-2 replication in AECs including human primary ALI-cultured AECs, facilitating cellular entry in a galectin-glycan interaction-dependent manner. Our transcriptomic data show that Gal-9 accelerates and exacerbates several virus-induced pro-inflammatory programs in AECs. These observations are highly relevant to the clinical manifestations and management of COVID-19, suggesting that circulating Gal-9 has a direct impact on viral infectivity and the cytokine milieu in the airway epithelium, which constitutes the critical initial site of SARS-CoV-2 attachment and infection (34). Collectively, our findings complement previous reports highlighting plasma Gal-9 level as a biomarker of severe COVID-19 disease, providing a novel molecular and immunologic framework linking Gal-9 activity to disease pathology (Extended Data Fig. 7). Importantly, our data build on a robust literature featuring Gal-9 as a key host factor regulating viral immunopathogenesis.

We observed potent Gal-9-mediated enhancement of SARS-CoV-2 cellular entry, using both pseudoviral constructs and wildtype replication-competent viruses. Previous studies have demonstrated that Gal-9 promotes HIV entry through retaining protein disulfide isomerase (PDI) on the CD4+ T cell surface (23). Therefore, we initially hypothesized that Gal-9 promoted SARS-CoV-2 entry by retaining or increasing viral receptor expression on the cell surface. However, our flow cytometry data revealed no impact of Gal-9 exposure on ACE2 or TMPRSS2 surface expression. These data were further validated by our transcriptomic analysis which failed to show any modulation of ACE2 or TMPRSS2 mRNA transcripts by Gal-9. Studies have found roles for Gal-9 in bridging pathogen glycans to host cell surface glycans to promote target cell attachment (35). We therefore conducted experiments to determine if this phenomenon underlies Gal-9-mediated enhancement of SARS-CoV-2 entry. Our data confirm the hypothesis that this enhancement is dependent on facilitating viral binding and attachment, presenting three pieces of evidence to support this conclusion: 1) the attachment of SARS-CoV-2 virions to the cell surface is enhanced by Gal-9 through galectin-glycan interactions. 2) Gal-9 directly enhances the binding of the viral spike protein to the ACE2 receptor. 3) Gal-9 effects on SARS-CoV-2 growth kinetics reflect facilitation of an immediate-early viral life cycle stage. These findings are compatible with a provocative hypothesis expressed by Arciniegas and colleagues (36) that multiple galectins act in concert through N-and O-linked glycans on the SARS-CoV-2 spike to form a galectin-glycan lattice on the virion surface that promotes viral attachment and penetration. Lujan and colleagues have recently shown that galectins including Gal-1, Gal-3, and Gal-9 can promote cell-surface binding, internalization and cell invasion of many sexually transmitted pathogens, including bacteria, parasites, and viruses (37). This enhancement is mediated by bridging the pathogen surface and receptors on host tissues in a glycosylation-dependent manner (37). Our data further revealed that the cellular entry of VSV-G, like SARS-CoV-2, is enhanced by Gal-9 exposure, suggesting that entry enhancement of multiple viral taxa could be mediated by galectin-glycan lattice interactions.

Accumulating evidence suggests that fatal COVID-19 is characterized by a profound cytokine storm (38). The overproduction of pro-inflammatory cytokines, such as IL-6, IL-8, TNFα, and IL-1β, leads to an increased risk of vascular hyperpermeability, acute lung injury, multiorgan failure, and eventually death when the high cytokine concentrations are unabated over time (39). In direct relation to this phenomenon, our transcriptomic analysis revealed that Gal-9 and SARS-CoV-2 infection synergistically induced the expression of key pro-inflammatory programs in AECs including the IL-8, IL-17, EIF2 and TNFα signaling pathways. These data were validated in part at the protein level using a bead-based immunoassay to characterize the AEC secretome. Bozorgmehr and colleagues demonstrated highly significant positive correlations of plasma Gal-9 levels with a wide range of pro-inflammatory biomarkers in COVID-19 patients. They further demonstrated that Gal-9 treatment of monocytes *in vitro* enhanced expression and production of key pro-inflammatory molecules associated with severe COVID-19 disease (6). Gal-9 induces secretion of inflammatory cytokines in several immune cell lineages including monocyte-derived macrophages and neutrophils (40, 41). Our findings validate and extend these observations to SARS-CoV-2 infection of the airway epithelium.

Importantly, the combined effects of SARS-CoV-2 infection and Gal-9 exposure on pro-inflammatory signaling are much stronger than either Gal-9 or SARS-CoV-2 alone, reflecting a synergistic interaction. We observed little transcriptional perturbation by SARS-CoV-2 alone after 24 hours of infection, which is consistent with other reports that profound transcriptional changes are only evident after 48 hours (42, 43). Interestingly, the IL-17 signaling pathway was significantly activated after Gal-9 treatment in the presence or absence of SARS-CoV-2. Our data are in accordance with a previous report demonstrating induction of IL-17 signaling by Gal-9 treatment *in vivo,* in the setting of sepsis (44). For MERS-CoV, SARS-CoV and now SARS-CoV-2, disease severity has been shown to positively correlate with levels of IL-17 and other T helper 17 (Th17) cell-related pro-inflammatory cytokines (45–47). IL-17 inhibition has been adopted as a common and successful strategy to reduce the injury associated with inflammatory autoimmune diseases (48). Thus, inhibition or neutralization of Gal-9 would not only decrease SARS-CoV-2 replication but also attenuate IL-17 signaling and other damaging pro-inflammatory cascades.

Our study has limitations that must be considered. Firstly, we focused exclusively on the airway epithelium, and it is now established that SARS-CoV-2 is capable of infecting other cell lineages including monocytes, monocyte-derived macrophages, and microglia (49–51). It is possible that Gal-9 does not exert similar effects on viral replication or immune signaling in other target cell types. Secondly, our studies were all performed *in vitro* or in transplant tissue-derived primary epithelial cells *ex vivo*. As it is well-established that Gal-9 exerts conditional, pleiotropic immunomodulatory effects, the net effect of Gal-9 signaling on SARS-CoV-2 pathogenesis cannot be fully appreciated outside of an animal model with a functional immune system. Validation and extension of our observations in murine, hamster or nonhuman primate models of SARS-CoV-2 infection will help in evaluating the clinical relevance of our reported findings. In relevance to the implementation of animal models, our study did not elucidate the principal cell or tissue sources responsible for secreting Gal-9 in the setting of SARS-CoV-2 infection. Leveraging an animal model to identify these source compartments will be critical in developing interventions to manipulate Gal-9 signaling as a therapeutic approach.

To our knowledge, the data presented here are the first to show that Gal-9 is directly involved in enhancing SARS-CoV-2 infection and virus-induced pro-inflammatory pathology, exposing an area of vulnerability for further investigation. Our data warrant examination of Gal-9-targeting therapeutic strategies *in vivo*, designed to simultaneously inhibit viral replication and suppress deleterious immune signaling associated with COVID-19 disease. There are existing tools which may be exploited to experimentally manipulate Gal-9 *in vivo*, including administration of exogenous, recombinant stable-form Gal-9 to enhance signaling (52). Small molecules and promising monoclonal antibodies under investigation in the clinical oncology space (e.g. LYT-200) that antagonize Gal-9 activity may have utility in COVID-19 (15, 53, 54). On this note, a recent exploratory clinical trial involved administration of a galectin antagonist to individuals with COVID-19 disease, revealing beneficial effects of galectin inhibition on disease outcomes (55). This represents a provocative and promising early step in the development of galectin-based therapeutics for SARS-CoV-2 infection in the future.

## Methods

### Cell lines

The Calu-3 human lung adenocarcinoma epithelial cell line was obtained from ATCC (ATCCHTB-55) and cultured in Eagle’s Minimum Essential Medium (EMEM). Vero E6 cells were purchased from ATCC (CRL-1586) and cultured in Dulbecco’s Modified Eagle’s medium (DMEM). Vero E6 cells stably expressing TMPRSS2 (Vero E6-TMPRSS2) were established and cultured in DMEM in the presence of puromycin (1µg/ml). All media were supplemented with 10% FBS and 1% penicillin/streptomycin. All cells had been previously tested for mycoplasma contamination and incubated at 37°C in a humidified atmosphere with 5% CO2.

### Primary airway epithelial cells (AECs)

Human bronchus was harvested from five explanted healthy lungs. The tissue was submerged and agitated for one minute in PBS with antibiotics and 5 mM dithiothreitol to wash and remove mucus. After three washes, the tissue was placed in DMEM with 0.1% protease and antibiotics overnight at 4°C. The next day, the solution was agitated and the remaining tissue removed. Cells were centrifuged at 300g/4°C for five minutes, then the cell pellet was resuspended in 0.05% trypsin-EDTA and incubated for five minutes at 37°C. The trypsinization reaction was neutralized with 10% FBS in DMEM, then cells were filtered through a cell strainer and centrifuged at 300g/4°C for five minutes. The cell pellet was resuspended in 10% FBS in DMEM and 10 μl was stained with trypan-blue and counted on a hemocytometer. 75,000 cells were plated onto each 6mm/0.4mm Transwell ALI insert after treatment with FNC coating mixture. 10% FBS in DMEM and ALI media were added in equal volumes to each basal compartment and cultures were incubated at 37°C with 5% CO_2_. The next day, the media was removed and both compartments were washed with PBS and antibiotics. ALI media was then added to each basal compartment and changed every three days until cells were ready for use at day 28.

### Viruses

The severe acute respiratory syndrome coronavirus 2 (SARS-CoV-2), isolate USA-WA1/2020 and SARS-Related Coronavirus 2 (SARS-CoV-2), isolate lineage P.1 were obtained from BEI Resources of National Institute of Allergy and Infectious Diseases (NIAID). Viruses were propagated in Vero E6-TMPRSS2 cells in DMEM with 2% FBS and 1% penicillin/streptomycin and viral stocks were stored at −80°C. Virus titer was measured in Vero E6 cells by TCID_50_ assay. All the studies involving live viruses were conducted in the Vitalant Research Institute BSL-3 under approved safety protocols.

### Cytotoxicity assay

The cytotoxic effect of Gal-9 on Calu-3 cells was determined using an MTT assay kit (abcam, ab211091) according to manufacturer’s guidelines. In brief, Calu-3 cells cultured in 96-well cell culture plates were incubated with different concentrations of Gal-9 (0-5,000 nM). After 48 h, the media was removed and 100□lμl MTT reagent (1:1 dilution in DMEM medium (serum free)) was added to each well and incubated for 3 h at 37°C. Then the medium was removed, and 150 μl MTT solvent was added into each well. Quantification was performed by reading absorbance at OD=590 nm. The data from three independent experiments was used to calculate the CC_50_ by nonlinear regression using GraphPad Prism 8.0 software.

### SARS-CoV-2 infection and Gal-9 administration

Stable-form recombinant Gal-9 was used in all experiments (20, 52). Calu-3 cells were seeded at 0.5 x 10^6^ cells per well in 0.5 ml volumes using a 24-well plate, or were seeded at 1 x 10^5^ cells per well in 0.1 ml volumes using a 96-well plate. The following day, cells were pretreated with or without Gal-9 for 6 h. Then viral inoculum (MOI of 0.01; 500 μl/well or 100μl/well) was prepared using EMEM containing indicated concentrations of Gal-9 and added to the wells. The inoculated plates were incubated at 37°C with 5% CO_2_. 24 h post infection, supernatants were collected and stored at - 80°C. Cells were lysed with TRizol (Thermo Fisher Scientific, 15596026) for RNA extraction or fixed with methanol : acetone (1:1) for IFA assay.

For infection of ALI cultured primary AECs, SARS-CoV-2 (diluted in ALI-culture medium, MOI=0.1) was added on to the apical chamber of inserts (250 μl) and the basal compartment (500 μl). Then the cultures were incubated for 2 hours at 37°C (5% CO_2_) to enable virus entry. Subsequently, the cells were washed and fresh ALI medium (500 μl) was added into the basal compartment. Cells were incubated at 37°C (5% CO_2_) and harvested for analysis at 36 hours post-infection.

### Viral titer by TCID_50_ assay

Viral production by infected cells was measured by quantifying TCID_50_. Vero E6 cells were plated in 96-well plates at 5 × 10^4^ cells per well. The next day, supernatants collected from Calu-3 cells were subjected to 10-fold serial dilutions (10^1^ to 10^11^) and inoculated onto Vero E6 cells. The cells were incubated at 37°C with 5% CO_2_. Three to five days post infection, each inoculated well was evaluated for presence or absence of viral CPE. TCID_50_ was calculated based on the method of Reed and Muench (56).

### Immunofluorescence microscopy

Cells were washed with 1 X PBS, then were fixed and permeabilized with cold methanol : acetone (1:1) for 10 min at 4°C. Next, cells were washed with 1 X PBS and incubated in a blocking buffer (5% goat serum (Seracare Life Sciences Inc, 55600007)) at room temperature for 30 min. Cells were then incubated with a primary antibody (monoclonal rabbit anti-SARS-CoV-2 nucleocapsid antibody (GeneTex, GTX135357)) in 1 X PBS (1:1,000) overnight at 4°C. The following day, cells were washed three times with 1 X PBS and incubated with a secondary antibody (Goat anti-Rabbit IgG (H+L) secondary antibody, FITC (Thermo Fisher, 65-6111)) in 1 X PBS (1:200) for 1 h at 37°C. Then cells were washed three times with 1 X PBS and incubated with DAPI (300 nM) (Thermo Fisher Scientific, D1306) for 5 min at room temperature. Images were acquired using a fluorescence microscope.

### RT-qPCR

Total RNA was extracted using the chloroform-isopropanol-ethanol method. 500 ng of RNA was reversed transcribed into cDNA in a 20 μl reaction volume using the RevertAid First Strand cDNA Synthesis Kit (Thermo Fisher Scientific, K1622) in accordance with the manufacturer’s instructions. RT-PCR was performed for each sample using Taqman Universal Master mix II, with UNG (Thermo Fisher Scientific, 4440038) or using PowerUp SYBR Green Master Mix (Thermo Fisher Scientific, A25780) on a ViiA7 Real time PCR system. Primers and probes for detection of the *RNaseP* gene and SARS-CoV-2 Nucleocapsid (*N*) gene were obtained from IDT (2019-nCoV RUO Kit (Integrated DNA Technologies, 10006713)). The expression level of the *N* gene was determined relative to the endogenous control of the cellular *RNaseP* gene. Primers for detection of *GAPDH*, *IL-6*, *IL-8*, and *TNF*_α_ were:

*GAPDH* forward: 5’-AGAAGGCTGGGGCTCATTTG-3’;
*GAPDH* reverse: 5’-AGGGGCCATCCACAGTCTTC-3’;
*IL-6* forward: 5’-GGAGACTTGCCTGGTGAAA-3’;
*IL-6* reverse: 5’-CTGGCTTGTTCCTCACTACTC-3’;
*IL-8* forward: 5’-ATGACTTCCAAGCTGGCCGTGGCT-3’;
*IL-8* reverse: 5’-TCTCAGCCCTCTTCAAAAACTTCTC-3’;
*TNF*_α_ forward: 5’-CCTCTTCTAATCAGCCCTCTG-3’;
*TNF*_α_ reverse: 5’-GAGGACCTGGGAGTAGATGAG-3’.

### RNA-seq and Ingenuity Pathway Analysis (IPA)

RNA concentration and quality was measured using High Sensitivity RNA ScreenTape Analysis (Agilent, 5067-1500). cDNA libraries were prepared using the Illumina TruSeq Stranded total RNA library prep kit (Illumina, 20020597) and sequencing was performed on the Illumina Nextseq 550 platform generating 75 bp paired-end reads. The quality of raw sequencing reads was assessed using FastQC. Differentially-expressed genes were identified by GSA or ANOVA in Partek® Flow® imported into the QIAGEN Ingenuity Pathway Analysis (IPA) software application. IPA was used to identify gene ontologies, pathways and regulatory networks to which differentially-expressed genes belonged to, as well as upstream regulators (57). Reads were also aligned to the SARS-CoV-2 isolate WA-1 and analyzed using the QIAGEN CLC Genomics Workbench.

### Pseudovirus production

VSVΔG-luciferase-based viruses, in which the glycoprotein (G) gene has been replaced with luciferase, were produced by transient transfection of viral glycoprotein expression plasmids (pCG SARS-CoV-2 Spike and pCAGGS VSV-G) or no glycoprotein control into HEK293T cells by TransIT-2020 as previously described (58). In brief, cells were seeded into 15-cm culture dishes and allowed to attach for 12 hours before transfection with 30 μg expression plasmid per plate. The transfection medium was changed at 18 hours post-transfection. The expression-enhancing reagent valproic acid was added to a final concentration of 3.75 nM, and the cells were incubated for three-four hours at 37°C with 5% CO_2_. Then cells were inoculated with VSVΔG-luc virus at a multiplicity of infection of 0.3 for 4 h before the medium was changed. 24 hours post infection, the supernatants were collected and filtered through a 0.45-μm syringe filter.

### Pseudovirus entry assay

Calu-3 cells were plated into 96-well plates. The following day, cells were pre-treated with indicated concentrations of Gal-9 for six hours. In order to block ACE2 on the cell surface, cells were pretreated with indicated concentrations of an anti-ACE2 antibody for one hour. An unrelated anti-goat IgG antibody was used as a control. Pseudovirus harboring either SARS-CoV-2 spike or VSV-G glycoprotein were diluted in EMEM containing indicated concentrations of Gal-9, and then were added to Calu-3 cells. Controls included wells with serum predetermined to possess neutralizing activity. Cells were incubated for 24 h at 37°C with 5% CO_2_. Supernatants were then removed, cells were lysed, and luciferase activity was read using a commercial substrate (Promega, E1500).

### Flow cytometry

Cells were detached with 10% EDTA containing Zombie NIR (1:300) (BioLegend, 423105) for 10 min at 37°C. Then cells were washed three times with 1 X PBS and incubated with human ACE2 Alexa Fluor 488-conjugated antibody (R&D Systems, FAB9332G) or human TMPRSS2 Alexa-Fluor 594-conjugated antibody (R&D Systems, FAB107231T) for 30 min at room temperature. Cells were washed three times with 1 X PBS again. Analytical flow cytometry was performed with BD LSRII flow cytometer. Data was analyzed using FlowJo.

### ELISA-based assessment of SARS-CoV-2 spike binding affinity to ACE2

Assessment of Gal-9 effects on the binding of the SARS-CoV-2 spike to human ACE2 was performed using a commercially available spike-ACE2 binding assay kit (CoviDrop SARS-CoV-2 Spike-ACE2 Binding Activity/Inhibition Assay Kit, EPIGENTEK, D-1005-48) following the protocol provided by the manufacturer. Gal-9 or positive inhibitor control was mixed with indicated amounts of recombinant human ACE2 protein, then added to an ELISA plate coated with recombinant SARS-CoV-2 spike protein and incubated at 37°C for 60 min. Unbound ACE2 was removed. The amount of captured ACE2, which is proportional to ACE2 binding activity, is then recognized by an ACE2 detection antibody and measured by reading the absorbance at a wavelength of 450 nm.

### Multiplex cytokine analysis

Cytokines in the cell culture supernatants were measured using the human cytokine storm 21-plex procartaplex panel human magnetic bead Luminex assay (Thermo Fisher, EPX210-15850-901), following the manufacturer’s instructions. Supernatants were subjected to 2% Triton X-100 solvent-detergent (SD) mix for virus inactivation prior to quantification. Assay standards and inactivated samples were incubated with fluorescent-coated magnetic beads pre-coated with respective capture antibodies in a 96-well black clear-bottom plate. After washing, biotinylated detection antibodies were incubated with cytokine-bound beads for 30 min. Finally, streptavidin-PE was added and incubated for another 30 min. Measurements were acquired using the Invitrogen Luminex 200 IS Reader. Data analyses were performed using Bio-Plex Manager™ 6.1.1 (Bio-Rad Laboratories, Hercules, CA, USA). Standard curves were generated with a 5-PL (5-parameter logistic) algorithm, reporting values for both MFI and concentration data.

### Image Analysis

To measure the frequency of infected cells, randomly-selected areas were imaged. Each treatment had three replicates. The percentage of GFP-positive cells was determined by dividing the number of GFP+ cells by the number of DAPI+ cells. All samples were analyzed at the same threshold values. For quantification of GFP+ cells, CellProfiler-3 software was used to determine the fraction of GFP+ cells. Briefly, we used the software pipeline CorrectIlluminationCalculate to calculate an illumination correction function for each of the two channels (DAPI (blue) and GFP (green)). We then used another pipeline, CorrectIlluminationApply, to load each image and correct its illumination using the pre-calculated functions. Next, we ran ColorToGray to change all slides to gray and ran IdentifyPrimaryObjects to identify the number of each channel. Finally, data were exported by using the ExportToSpreadsheet pipeline. The same threshold value was applied to the images of each area.

### Quantitative analysis of synergy

The combinatorial effects of Gal-9 treatment and SARS-CoV-2 infection on pro-inflammatory cytokine expression were analyzed using the SynergyFinder web application, implementing the Bliss Independence model. The Bliss model generates synergy scores from a response matrix (59).

### Statistical analysis

Statistical analysis was performed using GraphPad Prism version 8 software. Data were presented as means ± SEM or median. Data were analyzed for statistical significance using an unpaired or paired Student’s *t* test to compare two groups, or using a paired one-tailed Wilcoxon test. Only *p* values of 0.05 or lower were considered statistically significant (*p>0.05* [ns], *p*_≤_*0.05* [*], *p*_≤_ *0.01* [**], *p*_≤_*0.001* [***], *p*_≤_*0.0001* [****]).

## Acknowledgements

This study was supported by the Program for Breakthrough Biomedical Research, which is partially funded by the Sandler Foundation. Additional support was provided by National Institutes of Health grants R01MH112457 (SKP) and the University of California San Francisco-Gladstone Institute of Virology & Immunology Center for AIDS Research (P30 AI027763).

## Author Contributions

L.D. and S.P. initiated the project and designed the experiments. L.D. and M.B. performed MTT assay. L.D. and A.G. performed and processed RNA-seq. F.D. and J.G. isolated and prepared ALI-cultured primary airway epithelial cells (AECs). J.B. analyzed RNA-seq data. L.D., S.Y., and L.N. quantified GFP positive cells. L.D. and P.D. performed multiplex cytokine expression measurement. J.J. and G.S. performed SARS-CoV-2 stock. T.K. provided human recombinant Galectin-9. L.L and S.P. prepared the manuscript. Z.D., J.G., and S.P. jointly supervised the work.

## Extended Data

**Extended Data Fig. 1 Gal-9 promotes SARS-CoV-2 replication during the early stages of the viral life cycle.** (A) Schematic timeline of the pre-infection treatment and post-infection treatment experiments. In the pre-infection treatment experiments, Calu-3 cells were pre-treated with 250 nM Gal-9 for six hours. Cells were washed and incubated with SARS-CoV-2 (MOI=0.01) for one hour. Then cells were washed again and were supplemented with fresh media. 24 hpi, cells were harvested for RNA isolation and RT-qPCR targeting the *N* gene. In the post-infection treatment experiments, cells were infected with SARS-CoV-2 (MOI=0.01) for one hour and washed with PBS. Then cells were incubated with 250 nM Gal-9. After 24 h incubation, cells were harvested for RNA isolation and RT-qPCR targeting the *N* gene. (B) Virus production (measured as viral *N* gene expression) in Calu-3 cells in pre-infection and post-infection treatment scenarios. Copies of SARS-CoV-2 *N* were calculated using the 2^-ΔCt^ method and *RNaseP* threshold cycle (Ct) values were used for normalization. Data are representative of the results of three independent experiments (mean ± SEM). Statistical significance was analyzed by *t* test. *p*_≤_*0.05* [*], *p*_≤_ *0.01* [**], *p*_≤_*0.001* [***], *p*_≤_*0.0001* [****].

**Extended Data Fig. 2 Gal-9 treatment and SARS-CoV-2 infection induce secretion of select pro-inflammatory cytokines.** (A-D) Supernatants were harvested from Calu-3 cells with indicated treatments at 24 hours post-infection. Cytokine protein levels were determined by Luminex assay. Data are representative of the results of three independent experiments (mean ± SEM). Statistical significance was analyzed by *t* test. *p*_≤_*0.05* [*], *p*_≤_ *0.01* [**], *p*_≤_*0.001* [***], *p*_≤_*0.0001* [****].

**Extended Data Fig. 3 Synergistic effect of Gal-9 treatment and SARS-CoV-2 infection on expression of pro-inflammatory cytokines.** (A-C) Combinatorial induction of IL-6 (A), IL-8 (B), and TNFα (C) by Gal-9 treatment and SARS-CoV-2 infection are shown by the response matrix of relative expression. (D-F) The synergistic effects on IL-6 (D), IL-8 (E), and TNFα (F) of Gal-9 treatment and SARS-CoV-2 infection are shown by the 2D synergy landscape. When the synergy score is less than −10, the interaction between two treatments is likely to be antagonistic; from −10 to 10, the interaction between two treatments is likely to be additive; larger than 10, the interaction between two treatments is likely to be synergistic.

**Extended Data Fig. 4 Interactome depicting synergistic impact of Gal-9 treatment and SARS-CoV-2 infection on the canonical IL-17 signaling pathway.** Predictions were generated by IPA based on the differential expression of genes related to the IL-17 signaling pathway in Galectin-9 Infected vs Control Uninfected. Interactome symbols and color codes are defined in the embedded legend.

**Extended Data Fig. 5 Interactome depicting synergistic impact of Gal-9 treatment and SARS-CoV-2 infection on the canonical IL-6 signaling pathway.** Predictions were generated by IPA based on the differential expression of genes related to the IL-6 signaling pathway in Galectin-9 Infected vs Control Uninfected. Interactome symbols and color codes are defined in the embedded legend.

**Extended Data Fig. 6 Top enriched disease and functional pathways following Gal-9 treatment and SARS-CoV-2 infection.** Top enriched diseases and functions (as determined by IPA) in (A) Galectin-9 Uninfected vs Control Uninfected and (B) Galectin-9 Infected vs Control Uninfected. Diseases and functions were filtered using a FDR<0.05 cutoff with fc>2.

**Extended Data Fig. 7 Model of how Gal-9 enhances SARS-CoV-2 replication and inflammation in airway epithelial cells.**

