## Supplementary figures and images for "Human Galectin-9 Potently Enhances SARS-CoV-2 Replication and Inflammation in Airway Epithelial Cells"

### Extended Data Figure 1

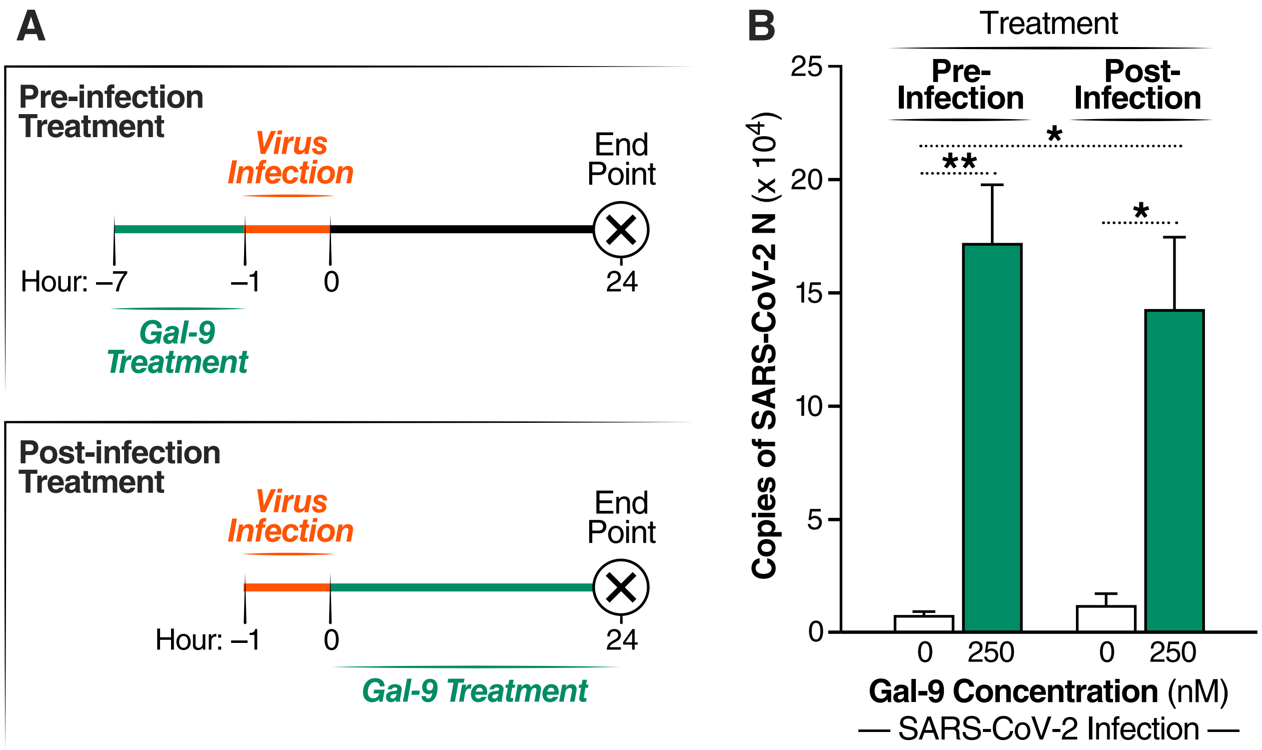

### Extended Data Figure 2

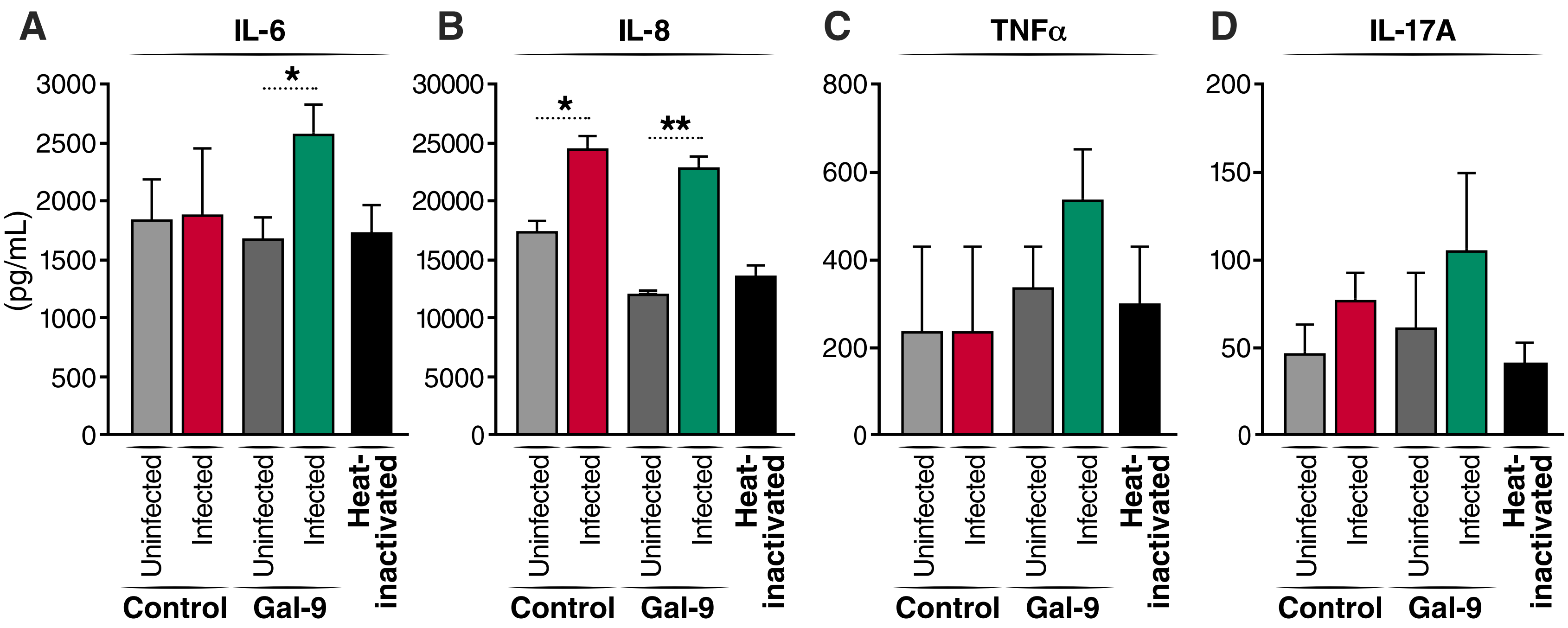

### Extended Data Figure 3

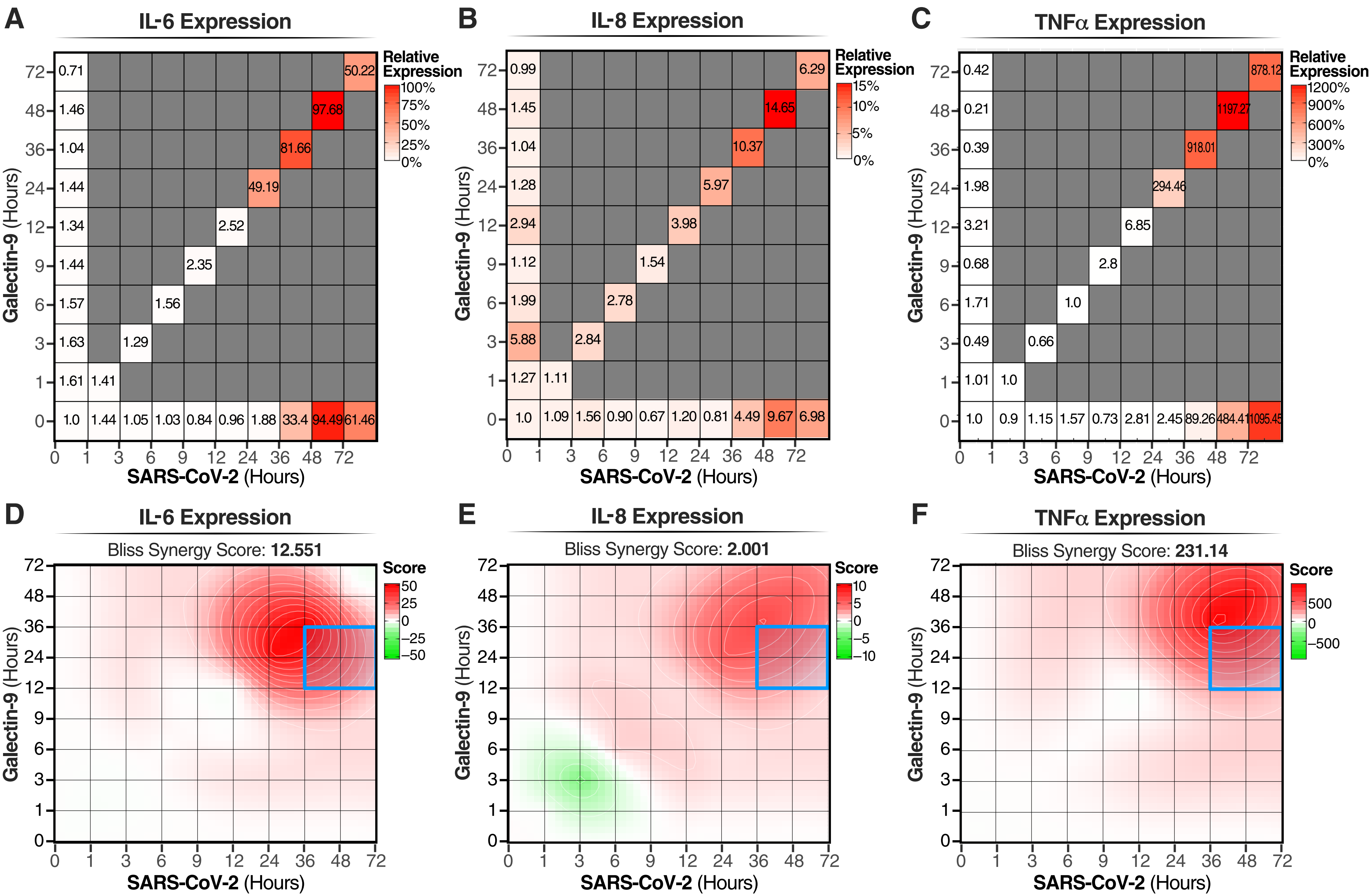

### Extended Data Figure 4

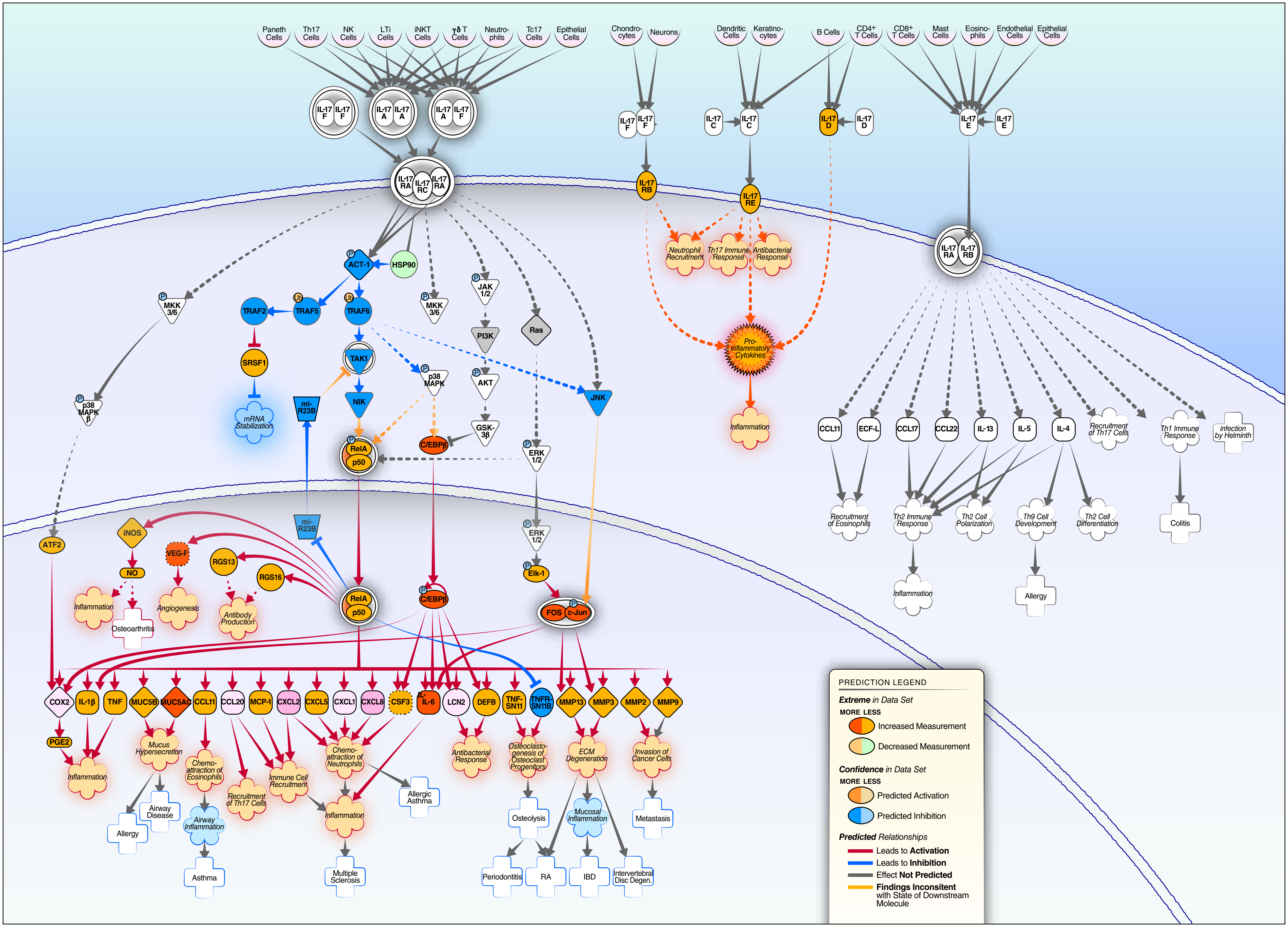

### Extended Data Figure 5

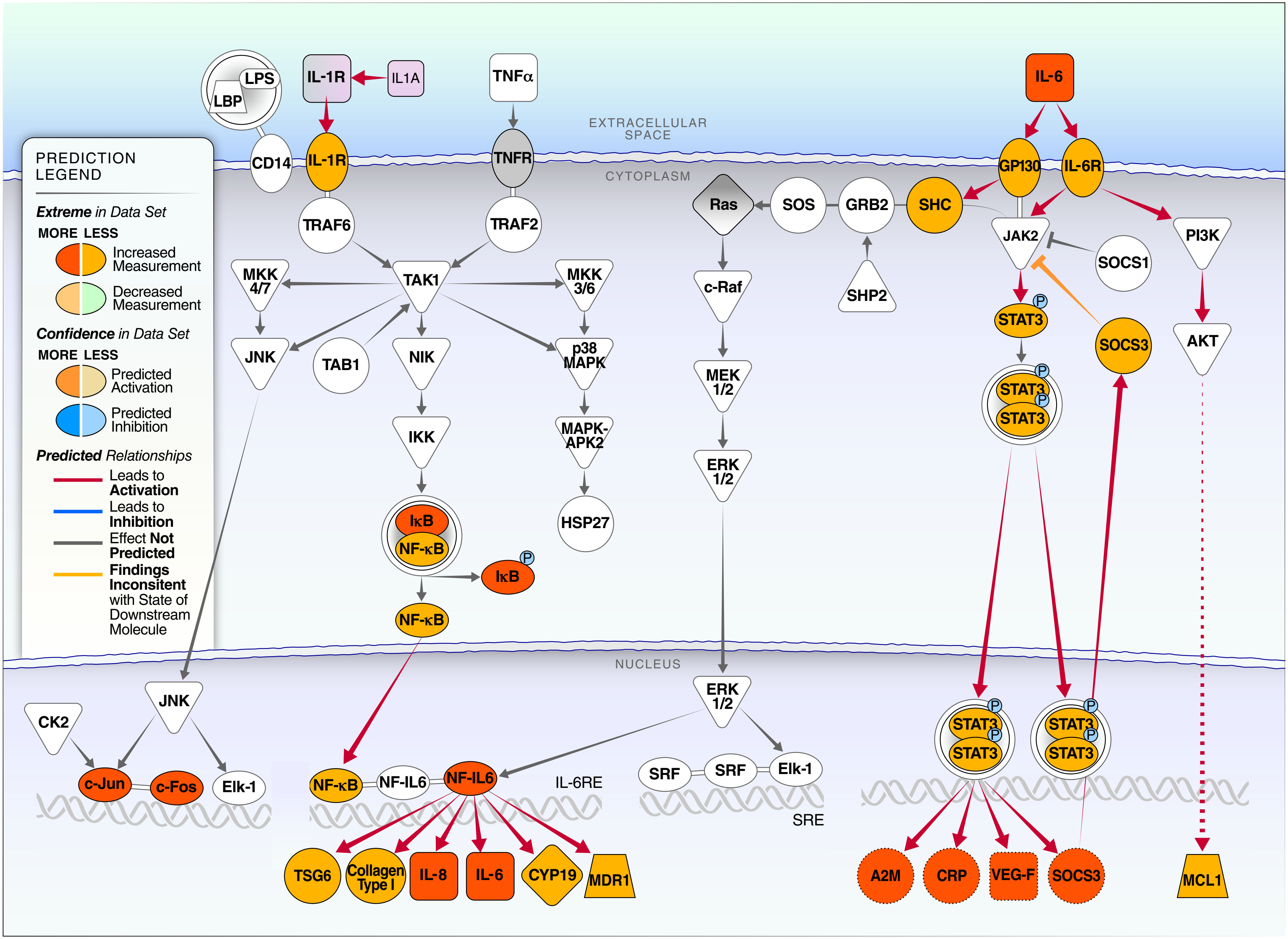

### Extended Data Figure 6

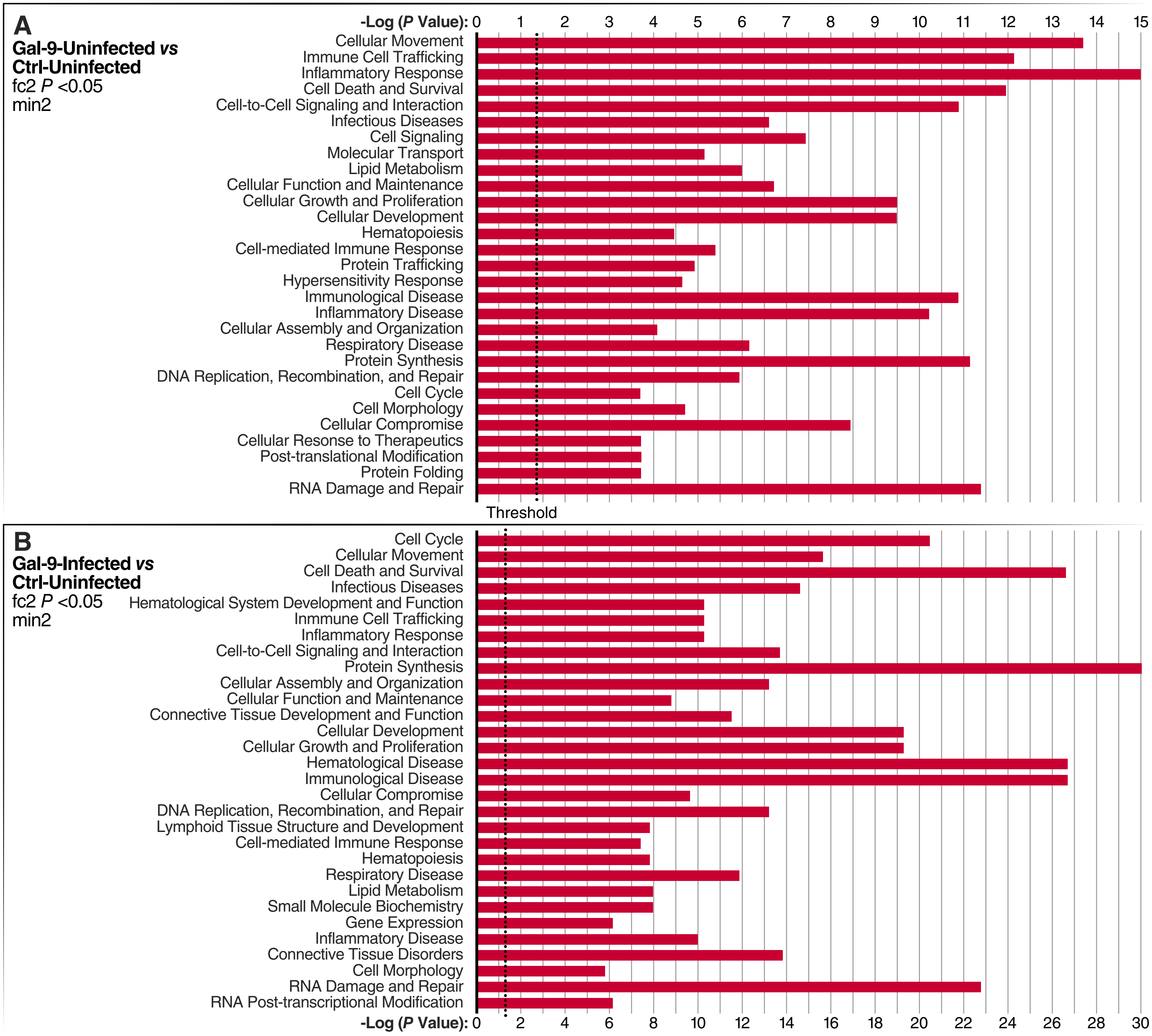

### Extended Data Figure 7

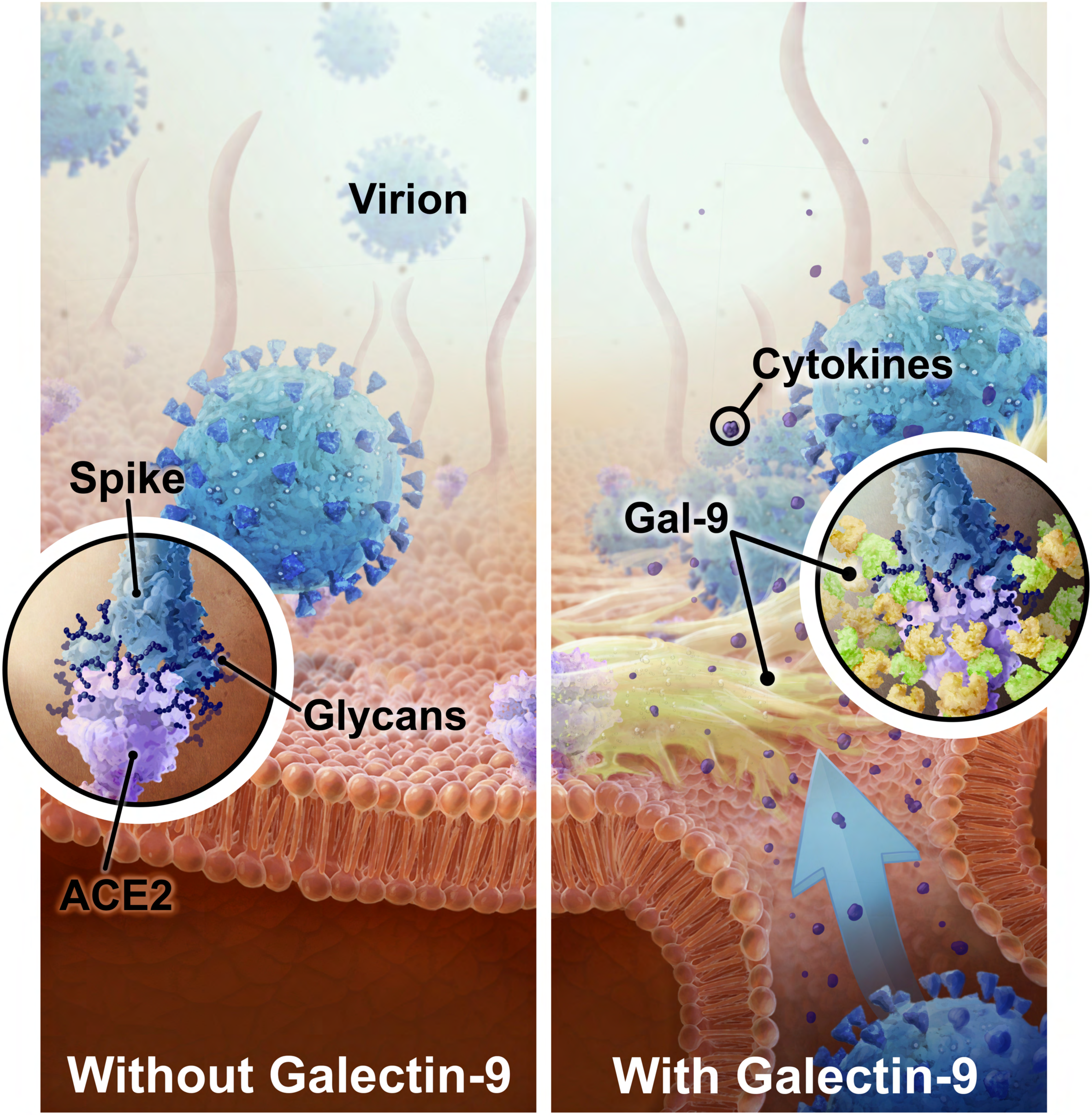
